# Quantifying the proportion of different cell types in the human cortex using DNA methylation profiles

**DOI:** 10.1101/2023.06.23.545974

**Authors:** Eilis Hannon, Emma L Dempster, Barry Chioza, Jonathan P Davies, Georgina ET Blake, Joe Burrage, Stefania Policicchio, Alice Franklin, Emma M Walker, Rosemary A Bamford, Leonard C Schalkwyk, Jonathan Mill

## Abstract

**Background:** Due to inter-individual variation in the cellular composition of the human cortex, it is essential that covariates that capture these differences are included in epigenome-wide association studies using bulk tissue. As experimentally derived cell counts are often unavailable, computational solutions have been adopted to estimate the proportion of different cell-types using DNA methylation data. Here, we validate and profile the use of an expanded reference DNA methylation dataset incorporating two neuronal- and three glial-cell subtypes for quantifying the cellular composition of the human cortex.

**Results:** We tested eight reference panels containing different combinations of neuronal- and glial-cell types and characterized their performance in deconvoluting cell proportions from computationally reconstructed or empirically-derived human cortex DNA methylation data. Our analyses demonstrate that these novel brain deconvolution models produce accurate estimates of cellular proportions from profiles generated on postnatal human cortex samples, they are not appropriate for the use in prenatal cortex or cerebellum tissue samples. Applying our models to an extensive collection of empirical datasets, we show that glial cells are twice as abundant as neuronal cells in the human cortex and identify significant associations between increased Alzheimer’s disease neuropathology and the proportion of specific cell types including a decrease in NeuNNeg/SOX10Neg nuclei and an increase of NeuNNeg/SOX10Pos nuclei.

**Conclusions:** Our novel deconvolution models produce accurate estimates for cell proportions in the human cortex. These models are available as a resource to the community enabling the control of cellular heterogeneity in epigenetic studies of brain disorders performed on bulk cortex tissue.

## Background

Recent years have seen acute interest in the role of epigenetic variation in the pathogenesis of disease. Although a number of different epigenetic mechanisms are involved in transcriptional regulation, the field of epigenetic epidemiology has focused primarily on DNA methylation (DNAm). DNAm can be quantified genome-wide using a commercial microarray(1, 2), making it cost effective to profile the large sample numbers required to detect statistically robust associations(3). Unlike genetic association studies, the choice of tissue for profiling epigenetic variation is a critical part of the study design for epigenome-wide association studies (EWAS). As the epigenome orchestrates the gene expression changes underpinning cellular differentiation, genome-wide patterns of DNAm are primarily defined by the tissue or cell type that the DNA sample originates from(4–7). Therefore, a major caveat of profiling DNAm in samples isolated from ‘bulk’ tissue, (e.g. whole blood or brain tissue) is that each is comprised of DNA from a heterogeneous mix of different cell types, with the resulting profile being an aggregate of each constituent cell type.

To date, most epigenetic datasets have been generated on DNA samples isolated from bulk tissues(8). As the proportion of each cell type within a sample can vary across individuals, systematic differences in cellular proportions that correlate with the phenotype of interest (e.g. pathology-associated changes in the abundance of a specific cell type) may manifest as differences in the overall epigenetic profile(9). For example, Alzheimer’s disease is characterised by extensive neuronal loss(10, 11) in conjunction with glial cell activation and proliferation in the cortex(12, 13). Adjusting analyses with quantitative covariates that capture the cellular composition of each sample has been widely adopted as the solution to minimising false positives. As experimentally derived cell counts are often not available, computational solutions have been proposed as an alternative. Deconvolution algorithms calculate a series of continuous variables reflecting the underlying cellular heterogeneity of each sample from the bulk tissue profile. Deconvolution algorithms can be separated into two classes - supervised methods (known as ‘reference based’ algorithms)(14–20) and unsupervised methods (known as ‘reference free’)(21–24)

Reference based methods in particular have been successfully used to control for cellular heterogeneity in DNAm studies of whole blood(25). However, because this approach requires reference DNAm profiles for each constituent cell type of interest, they are not applicable to the study of all tissues. Similarly, although reference profiles exist for deconvoluting cellular proportions from DNAm data generated on bulk cortex tissue, these are currently limited to estimating the abundance of neuronal and non-neuronal cells(16) - and do not capture the full complexity or diversity of cell types present in the brain(26, 27). We and others have recently developed experimental protocols using Fluorescence-Activated Nuclei Sorting (FANS) to purify populations of nuclei from multiple cell types in post-mortem human cortex tissue(28–30). These methods have enabled us to refine the non-neuronal (predominantly glial) cell population and generate reference DNAm profiles for oligodendrocyte, microglia and astrocyte nuclei that can be used for the cellular deconvolution of DNAm data generated on bulk cortex.

In this study, we profile the use of these novel cell reference datasets in conjunction with the widely used Houseman deconvolution algorithm(15) - a constrained projection methodology - for quantifying the cellular composition of the human cortex. First, we validate the use of these reference data with computationally simulated of ‘bulk’ cortex profiles, where the proportion of different cell-types is predetermined. Second, we apply these reference panels to empirical DNAm datasets generated from bulk cortex tissue samples to profile how deconvolution performance, as well as cellular composition, varies across brain regions and development. Finally, we demonstrate how the quantification of these refined brain cell types can be used as phenotypic variables for detecting known cellular changes associated with neuropathology in Alzheimer’s disease. To enable the wider research community to incorporate our novel cellular composition estimates into their workflow, our enhanced reference panels are available via the R CETYGO package on GitHub (https://github.com/ds420/CETYGO). Beyond the estimation of cell-type proportions in the human cortex, our analyses provide broader insights into the methodology of cellular deconvolution that are applicable for studies involving other cell types and tissues.

## Results

### Further refinement of neural cell types confirmed with distinct genome-wide DNAm profiles

We used a FANS protocol previously described by our group(31) to purify nuclei populations from prefrontal cortex tissue dissected from 43 adult donors. Our initial gating strategy used an antibody against NeuN (a robust marker of post-mitotic neurons (32)) to isolate neuronal nuclei in combination with an antibody against SOX10 (a transcription factor involved in the differentiation of oligodendrocytes(33)) to distinguish oligodendrocyte nuclei from other glial nuclei (**Supplementary Figure 1A**). Subsequently, in a second gating strategy we additionally included an antibody against IRF8 (a transcription factor that is upregulated in microglia(34)) to enrich microglia from the NeuNNeg/SOX10Neg fraction (**Supplementary Figure 1B**). Our third gating strategy used an antibody against SATB2 (a DNA binding protein involved in transcriptional regulation and chromatin remodelling which is expressed in excitatory neurons in the mature central nervous system(35)) in place of NeuN (**Supplementary Figure 1C**). We generated DNAm profiles using the Illumina EPIC array for NeuNPos (neuron-enriched; n = 28), NeuNNeg/SOX10Pos (oligodendrocyte-enriched; n = 24), NeuNNeg/SOX10Neg (microglia- and astrocyte-enriched; n = 21), NeuNNeg/SOX10Neg/IRF8Pos (microglia-enriched; n = 17), NeuNNeg/SOX10Neg/IRF8Neg, (astrocyte-enriched; n = 7), SATB2Pos (excitatory neuron-enriched; n = 9), and SATB2Neg (inhibitory neuron- and glial-enriched; n = 6) nuclei populations (**Supplementary Tables 1** & **2**). To confirm that cell type differences were the primary drivers of variation in DNAm across samples, principal component (PC) analysis was used (**Supplementary Figure 2**). The first PC, which explains 43.2% of the variance in DNAm, separates the NeuNPos fractions (NeuNPos and SATB2Pos) from the other nuclei populations. The second PC, which explains 28.8% of the variance, separates the NeuNNeg/SOX10Neg/IRF8Pos samples from the NeuNNeg/SOX10Neg/IRF8Neg samples, with NeuNNeg/SOX10Neg samples, the parent fraction, in between these extremes. While the third PC, which explains, 3.7% of the variance, does highlight differences between nuclei fractions, this does not correlate with any of the antibodies we used to isolate specific cell types. It appears to capture a difference between the NeuNNeg/SOX10Neg and the NeuNNeg/SOX10Neg/IRF8Pos fractions with NeuNNeg/SOX10Neg/IRF8Neg sitting in the middle. This could indicate that there is another cell type, which we have not isolated, characterised as NeuNNeg/SOX10Neg/IRF8Neg that is lost during the IRF8 gating but retained in the NeuNNeg/SOX10Neg fraction. All subsequent PCs, which each explain <3% of the variance, do not correlate with a specific nuclei population and therefore likely reflect technical or biological sources of variation in DNAm between samples.

In order to increase the specificity of brain cell types in our subsequent deconvolution analyses, we augmented our data with publicly available data from the EpiGABA(36) study in which the NeuNPos nuclei population is further refined using an antibody against SOX6(37) (**Supplementary Figure 1D**) using the Illumina 450K array to generate NeuNPos/SOX6Pos (GABAergic neuronal enriched; n = 4), NeuNPos/SOX6Neg (glutamatergic neuronal enriched; n = 3) and NeuNNeg (glial enriched; n = 4) nuclei populations isolated from occipital cortex tissue. PC analysis of this combined dataset (123 samples from 47 donors; **Figure 1**) showed that PC1 (explaining 39.9% of the variance) still separates neuronal and non-neuronal nuclei, with the NeuNPos/SOX6Pos and NeuNPos/SOX6Neg clustering with the NeuNPos and SATB2Pos samples and the NeuNNeg clustering with the other glial fractions. PC2 (explaining 23.9% of the variance) still separates NeuNNeg/SOX10Pos from NeuNNeg/SOX10Neg/IRF8Pos, with NeuNNeg samples located in-between these extremes reflecting the fact that this population contains nuclei from both of these subfractions. PC3 (explaining 11.7% of the variance) separates the two sets of data and likely reflects technical differences (e.g. different array types and other experimental batch effects). These results highlight that the major cell type differences in DNAm are highly reproducible across data generated in different laboratories and dominate over batch effects and inter-individual differences. We therefore decided that for the purposes of generating the most extensive set of cellular composition estimates we would merge our data with the EpiGABA DNAm data into a single dataset.

**Figure 1.**
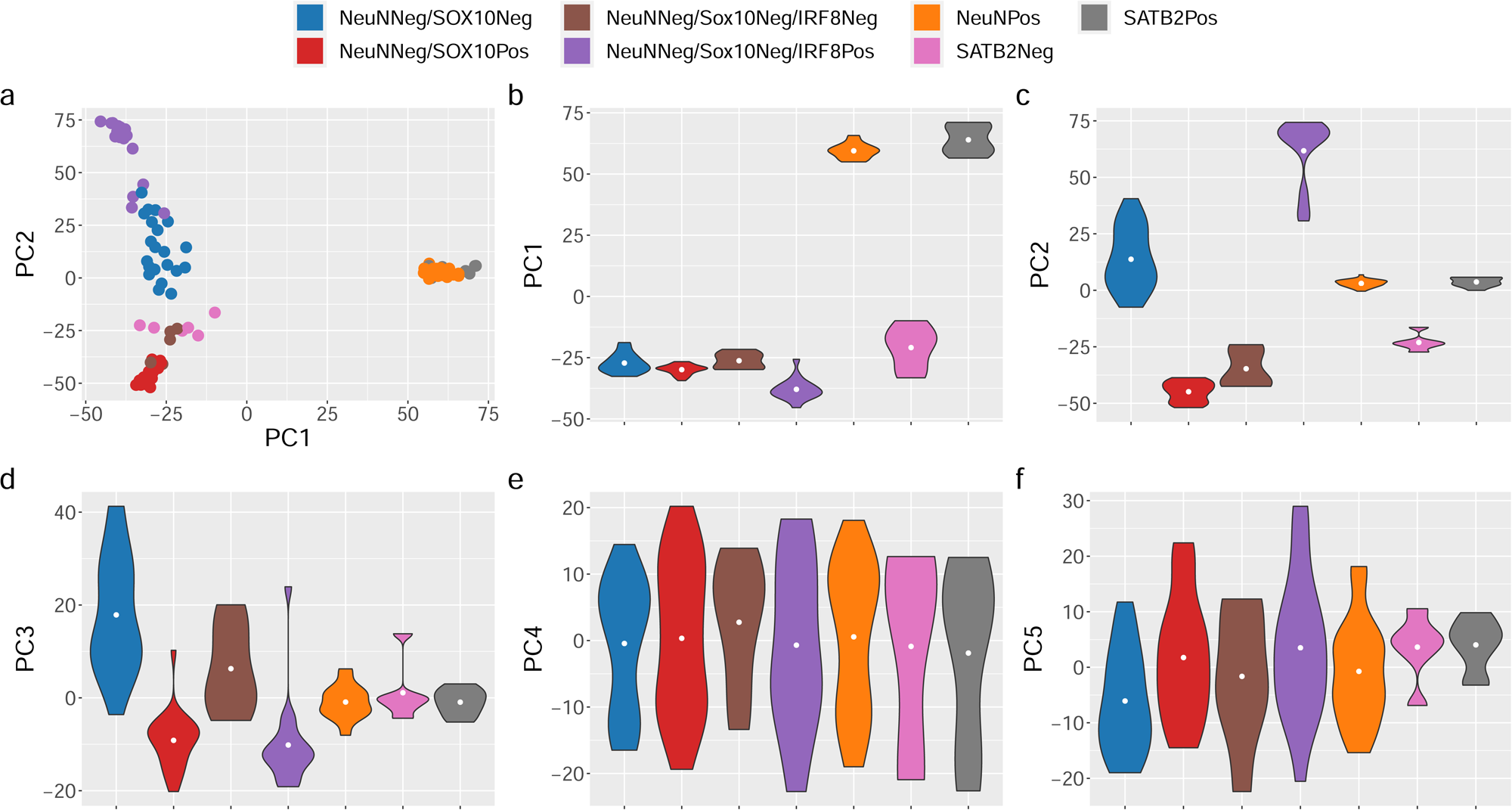
Major axes of variation in DNA methylation data are driven by cell type. Scatterplot (a) of the first two principal components where each point represents a sample and the colour of the point indicates the nuclei fraction. Violin plots for the first 5 principal components (b-f) grouped by nuclei fraction.

### Accuracy of cellular composition estimation depends on the combination of cell types included in the reference panel

Given the large number of nuclei fractions included in our final DNAm reference dataset, some of which target overlapping cell populations due to the different FANS gating strategies used, we defined 8 different combinations of cell types to serve as reference panels for the deconvolution of cellular composition of cortical DNAm data (**Table 1**, **Supplementary Figure 3**). Six of these represent mostly complete, non-overlapping and increasingly refined combinations, whereby any given cell type should be contained within a single fraction. These enabled us to characterise how deconvolution performance was affected by increasing the specificity of cellular composition. Two of the panels (4 and 5), contain overlapping fractions (SATB2Pos and NeuNPos), that both capture excitatory neuronal nuclei. These panels were included to observe how the algorithm handles this direct conflict.

**Table 1.** Summary of fractions included in the reference panels

| Panel | Included Fractions |
| --- | --- |
| 1 | NeuNPos<br>NeuNNeg/SOX10Pos<br>NeuNNeg/SOX10Neg |
| 2 | NeuNPos<br>NeuNNeg/SOX10Pos<br>NeuNNeg/SOX10Neg/IRF8Pos<br>NeuNNeg/SOX10Neg/IRF8Neg |
| 3 | SATB2Pos<br>SATB2Neg |
| 4 | NeuNPos<br>SATB2Pos<br>NeuNNeg/SOX10Pos<br>NeuNNeg/SOX10Neg |
| 5 | NeuNPos<br>SATB2Pos<br>NeuNNeg/SOX10Pos<br>NeuNNeg/SOX10Neg/IRF8Pos<br>NeuNNeg/SOX10Neg/IRF8Neg |
| 6 | NeuNPos/SOX6Pos<br>NeuNPos/SOX6Neg<br>NeuNNeg |
| 7 | NeuNPos/SOX6Pos<br>NeuNPos/SOX6Neg<br>NeuNNeg/SOX10Pos<br>NeuNNeg/SOX10Neg |
| 8 | NeuNPos/SOX6Pos<br>NeuNPos/SOX6Neg<br>NeuNNeg/SOX10Pos<br>NeuNNeg/SOX10Neg/IRF8Pos<br>NeuNNeg/SOX10Neg/IRF8Neg |

To compare the performance of the different panels, we performed a series of simulations where we could contrast predicted composition against a known truth. Briefly for each panel, we held one sample of each nuclei fraction back, selected the sites for deconvolution using all other samples for that fraction. We then used the excluded sample to construct bulk brain DNAm profiles where we combined cell-specific profiles in a weighted linear sum of pre-specified proportions of each cell type (see **Methods**). As well as comparing 8 different combinations of cell types, for panels with > 2 fractions, we also compared two methods, ANOVA and IDOL (IDentifying Optimal Libraries) algorithm(19), for selecting cell-specific sites that are the basis of the algorithm. In total 15 different training models were considered in the Houseman constraint projection deconvolution methodology(15) using these learnt parameters to estimate the cellular composition of a bulk profile. Overall accuracy of the deconvolution was captured by two metrics, the CETYGO score(38), which quantifies the accuracy of cellular deconvolution where the true cellular composition is unknown, and root mean square error (RMSE), which requires the cellular composition to be known.

In general, each reference panel combination yielded highly accurate estimates of cell proportions (average CETYGO < 0.10 using either ANOVA or IDOL) with performance being comparable across the different panels and site selection methods (**Figure 2**, **Supplementary Figure 4, Supplementary Table 3**). For each reference panel, we performed the deconvolutions with increasing numbers of cell-specific sites but found that this had little effect on the accuracy of the deconvolution (**Supplementary Figure 5, Supplementary Figure 6**). Marginally the best panel, measured by both the CETYGO score and RMSE was panel 6 (NeuNPos/SOX6Pos, NeuNPos/SOX6Neg, NeuNNeg). Of note, the separation of the NeuNNeg/SOX10Neg fraction into NeuNNeg/SOX10Neg/IRF8Pos and NeuNNeg/SOX10Neg/IRF8Neg (e.g. comparing panel 1 with panel 2) was associated with a slightly lower CETYGO score, indicative of a composition profile that captured more of the variation in the bulk tissue. This was generally also true of the separation of the NeuNPos fraction into NeuNPos/SOX6Pos and NeuNPos/SOX6Neg fractions (e.g. comparing panel 2 with panel 8) although not ubiquitously the case. In contrast, more refined cellular deconvolution models (i.e. incorporating more cell types) were associated with a slightly higher RMSE (**Supplementary Figure 4**) indicating that although the inclusion of more cell types gives a better representation of the variation in a bulk tissue, the estimates of the individual fractions are associated with a higher degree of error. We also observed this pattern when comparing the reference panels that consist of both SATB2Pos and NeuNPos (panels 4 and 5).

**Figure 2.**
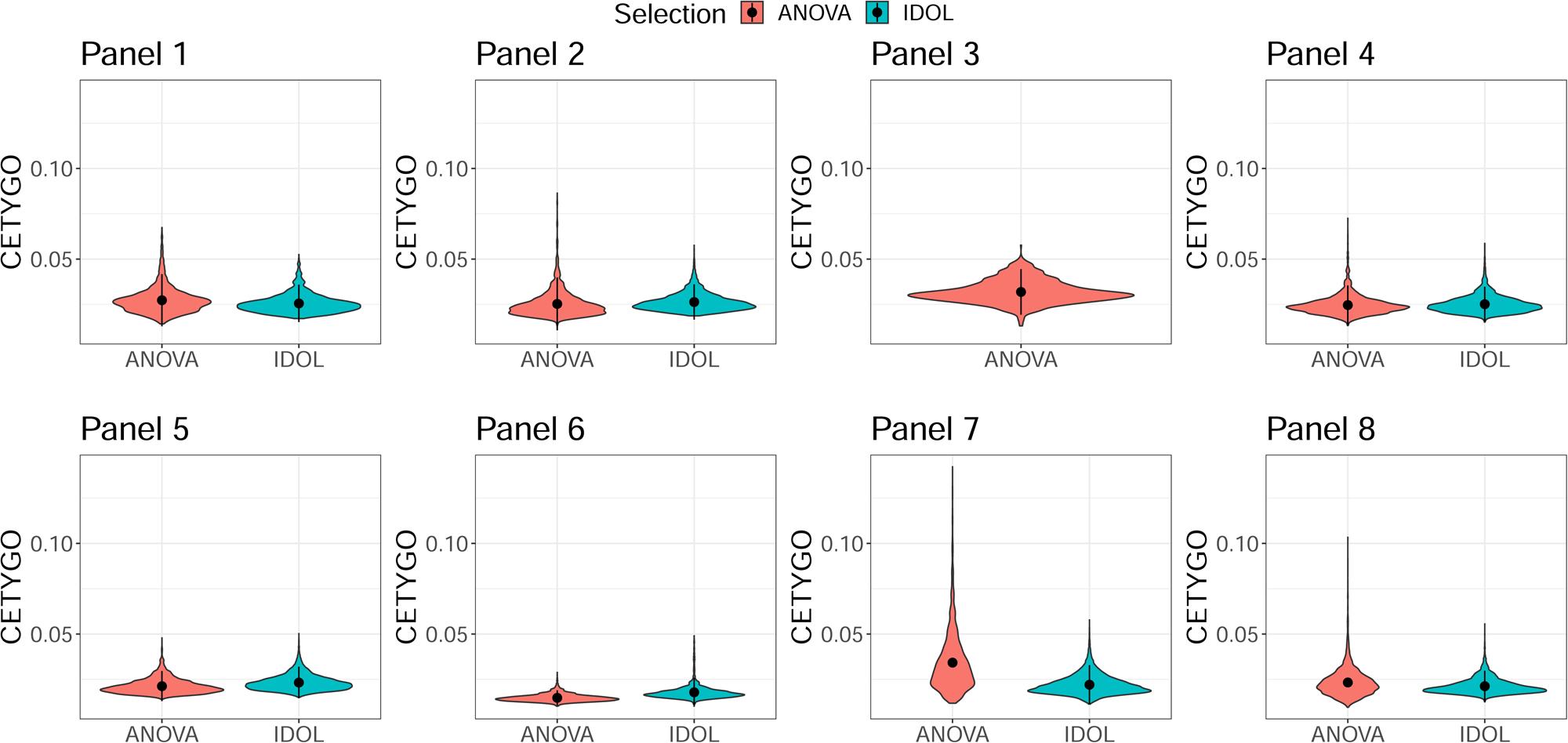
Accurate and increasingly refined estimation of cellular composition of the cortex from DNA methylation profiles. Violin plots of the error, measured by CETYGO, associated with estimating the cellular proportions of reconstructed cortical brain profiles. Panels represent different combinations of nuclei populations, as defined in **Table 1**. For reference panels with more than two cell types, two methods were used to select the cell-specific sites that serve as the basis for the algorithm represented by different violins.

Looking more specifically at the accuracy of estimating the proportion of particular nuclei fractions, we observe noticeable variation in the degree of accuracy (**Figure 3**, **Supplementary Table 4, Supplementary Figures 7-14**). Some cell types performed consistently accurately, regardless of which reference panel was used. Furthermore, we could group cell types based on their summary statistics. As described above, the accuracy of estimating the proportion of NeuNPos and SATB2Pos nuclei was dramatically reduced in the two reference panels (4 & 5) where they were both included and therefore all subsequent analyses focused on panels were either one or the other was used. The top performing cell fractions with near perfect estimates included NeuNPos, NeuNNeg, NeuNPos/SOX6Pos and NeuNPos/SOX6Neg (all r ≥ 0.99 and RMSE ≤ 0.02, **Supplementary Table 4**). NeuNNeg/SOX10Neg, NeuNNeg/SOX10Neg/IRF8Pos, SATB2Pos, and SATB2Neg are associated with marginally larger errors but still perform well with r ≥ 0.92 and RMSE ≤ 0.06. Of note the NeuNNeg/SOX10Pos fraction showed the most variation across panels. When included in a panel where the NeuNNeg/SOX10Neg fraction was replaced with the NeuNNeg/SOX10Neg/IRF8Pos and NeuNNeg/SOX10Neg/IRF8Neg fractions, this had a dramatic effect on the accuracy of NeuNNeg/SOX10Pos estimates, with the correlation statistic (r) decreasing from ∼ 0.95 to ∼ 0.7 and the RMSE doubling from ∼ 0.05 to > 0.1. The best statistics for predicting the NeuNNeg/SOX10Neg/IRF8Neg fraction come from panel 5 (which interesting includes both SATB2Pos and NeuNPos) with r = 0.81 and RMSE = 0.09; of note this fraction provides the least accurate prediction metrics. Instead considering the (signed) error, we observed that some cell types were associated with a particular bias in their estimation; for example, both NeuNNeg/SOX10Neg (median error = 0 - 0.022) and NeuNNeg/SOX10Neg/IRF8Pos (median error = 0.007 – 0.025) were typically overestimated (**Figure 3, Supplementary Table 4**). These results highlight how the accuracy of prediction for a given cell type is influenced by which other cell types are included in the deconvolution model, even when using a non-overlapping reference panel. Additionally, our results indicate that the accurate estimation of one cell type in a panel does not necessarily mean that the proportions of other cell types in that panel are also well estimated. A natural consequence of these conclusions is that to get the most precise estimates of a diverse set of cell types, different reference panels may need to be utilised in parallel. All these analyses were repeated using the IDOL method for selecting cell-specific sites for deconvolution, and there was no clear evidence that one method for selecting cell-specific prediction sites outperformed the other (**Figure 3**, **Supplementary Figure 4**).

**Figure 3.**
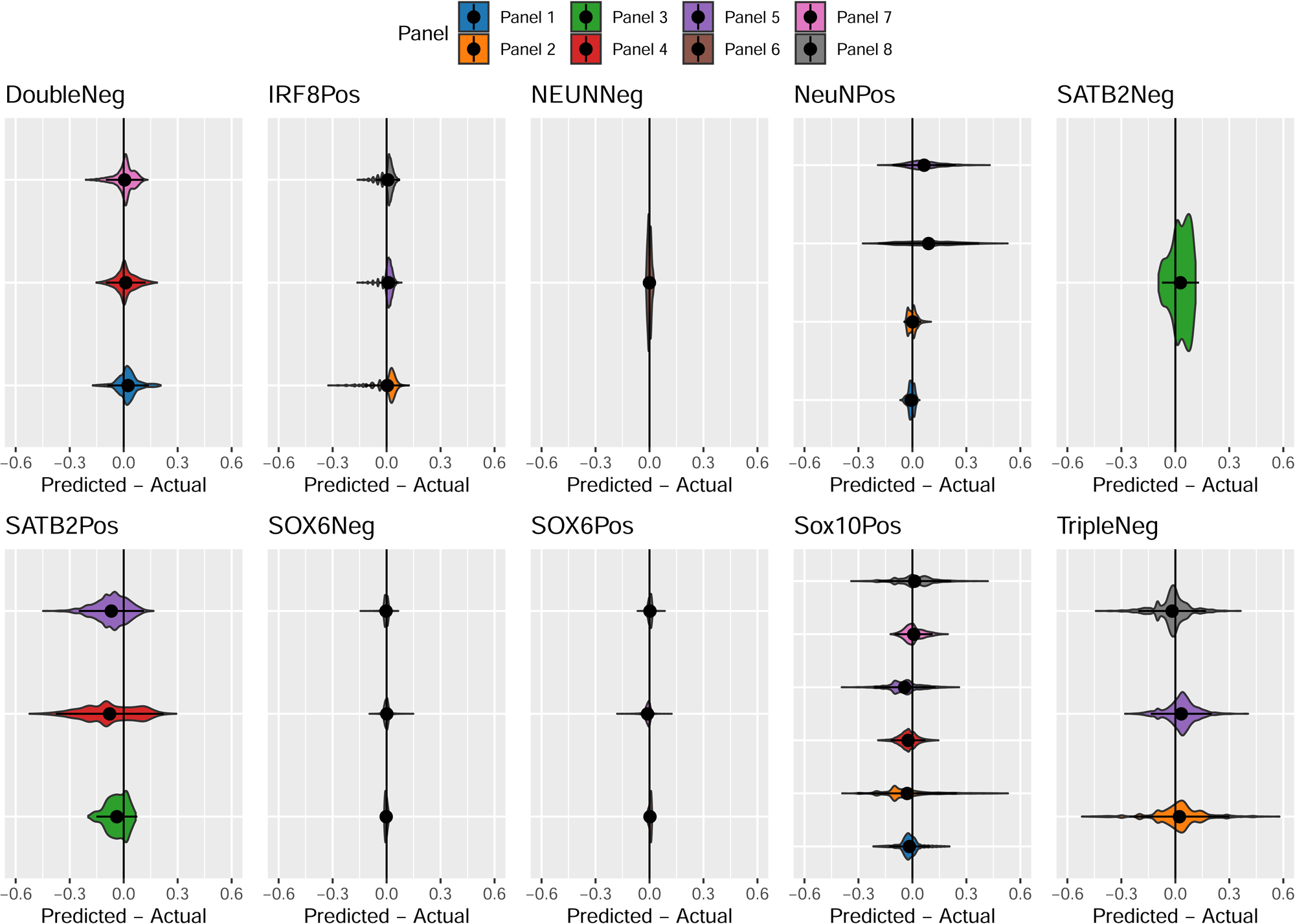
Accuracy and bias differs across cell types when estimating the cellular composition of the cortex. Violin plots of the error associated with estimating the cellular proportions of reconstructed cortical brain profiles, measured as the difference between predicted and actual abundance, where a positive value indicates an overestimation. Panels collate results for the same cell type, within each panel, values are grouped by reference panels, as defined in **Table 1**.

### Technical variation influences the accuracy of cellular deconvolution

Having demonstrated that our new reference panels for cellular deconvolution are capable of calculating accurate estimates of cellular composition in the cortex, we used them to calculate estimates in two large bulk DNAm datasets generated using adult prefrontal cortex tissue. The first dataset (the ‘Exeter’ dataset) incorporates a number of datasets generated by our group (http://www.epigenomicslab.com) (n = 377, age range = 19-108 years old)(39–43) and the second represents a publicly available dataset described by Jaffe *et al.* (n = 415, age range = 18-97 years old) (44). Profiling the accuracy of the deconvolution using the CETYGO score highlighted that all panels performed well (mean CETYGO < 0.10), with reference panel 6 (NeuNPos/SOX6Pos, NeuNPos/SOX6Neg, NeuNNeg) being associated with the lowest scores (**Supplementary Figure 15**) consistent with the simulation results. This was closely followed by panels 7 (NeuNPos/SOX6Pos, NeuNPos/SOX6Neg, NeuNNeg/SOX10Pos, NeuNNeg/SOX10Neg) and 8 (NeuNPos/SOX6Pos, NeuNPos/SOX6Neg, NeuNNeg/SOX10Pos, NeuNNeg/SOX10Neg/IRF8Pos, NeuNNeg/SOX10Neg/IRF8Neg), with the other 5 panels performing similarly. Of note, CETYGO scores were strongly correlated across panels (**Supplementary Figure 16**), suggesting that regardless of reference panel, there are other important influences on the accuracy of the estimates, such as data quality.

Subsequently, testing for biological or technical factors that influence the accuracy of cellular deconvolution we found that the CETYGO score was significantly associated with batch (**Figure 4A)** in both datasets, across all reference panels (**Supplementary Table 5**). There was a significant effect (P < 0.00333 corrected for 15 training models) of sex on the CETYGO score for 4 models in the Exeter dataset and 10 models in the Jaffe dataset (**Supplementary Table 5**). In all cases, females were associated with a slightly lower average error (**Supplementary Figure 17**) especially when the ANOVA method was used to select cell-specific sites (11/14 significant associations), despite more male samples being included. Of note, there was no association with age or age squared on prediction accuracy in either dataset (**Supplementary Table 5**).

**Figure 4.**
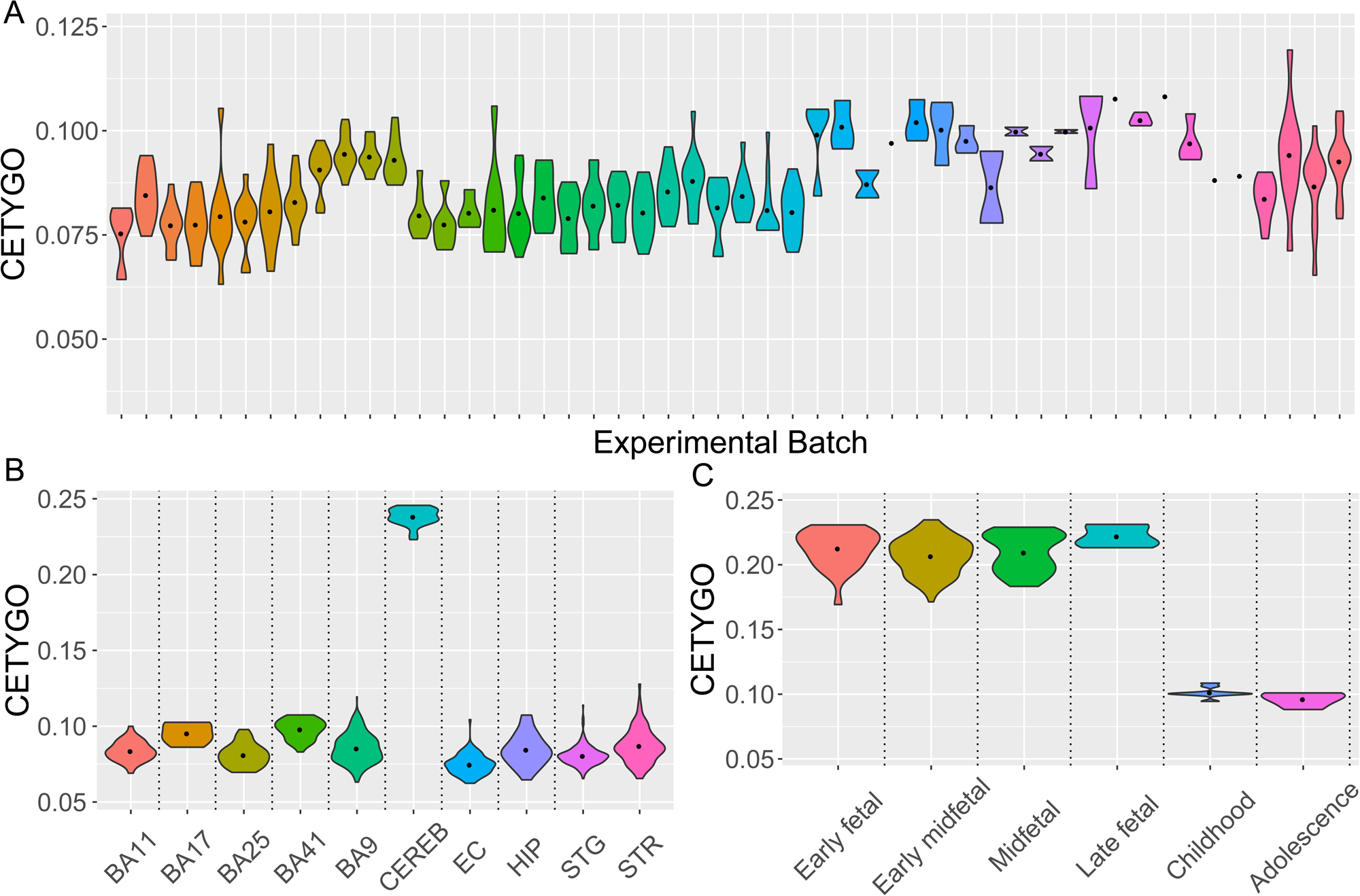
Performance of brain cellular deconvolution models not equitable across all brain datasets. Violin plots of the distribution of the error, measured by CETYGO, associated with estimating the cellular proportions from DNA methylation data generated from brain tissues. **A**) Adult prefrontal cortex samples grouped by experimental batch (n = 377). **B**) Adult brain samples grouped by brain region (n = 851). **C**) Prenatal and childhood cortical samples grouped by developmental stage (n = 167). CETYGO scores taken from reference panel 1 with an ANOVA to select cell specific sites.

### Neural cellular deconvolution panels derived from adult cortical samples do not effectively capture cellular heterogeneity in the cerebellum or fetal DNAm datasets

While our reference profiles were generated from populations of nuclei isolated from prefrontal and occipital cortical tissue, they are potentially relevant for estimating the proportion of the same cell types in other brain regions, especially other regions of the cortex. We performed cellular deconvolution using DNAm profiles from an additional 851 samples (age range = 19 -108 years old)(39-42, 45, 46) generated by our group from 9 other brain regions including additional cortical regions, the striatum, the hippocampus, and cerebellum (**Supplementary Table 6**). These analyses showed that the CETYGO scores in cerebellum samples are dramatically elevated, indicating that the cellular composition estimates for this tissue are unlikely to be accurate (**Figure 4B**, **Supplementary Figure 18**). It is known, for example, that the predominant neuronal subtype in the cerebellum (Purkinje cells) do not express NeuN(47). We also observe subtle differences in performance between the other 8 regions, although the distribution of CETYGO scores largely overlap with those observed in the prefrontal cortex (**Supplementary Table 7**).

We also wanted to confirm whether our reference panels were suitable for use in samples from donors at earlier stages of development. To this end we used 167 prenatal and childhood DNAm profiles generated from bulk cortex samples by our group (age range = 23 days post conception – 17 years old)(39, 48). We found consistently elevated CETYGO scores in the prenatal samples regardless of the specific developmental stage, comparable with those seen in the cerebellum samples (**Figure 4C**, **Supplementary Figure 19**) suggesting that bespoke reference panels are required to estimate cellular proportions in prenatal cortex tissues. The distribution of postnatal childhood and adolescent samples CETYGO scores are comparable to adult scores. Of interest, reference panel 3 has the smallest difference between prenatal and postnatal CETYGO scores reflecting the fact that SATB2 is a more robust marker of neuronal cells than NeuN in the prenatal cortex(49).

### Variable abundance of neuronal and glial cells in the adult prefrontal cortex

While there has been a fair degree of interest in profiling the cellular heterogeneity of the brain, variation in study design and methodologies have made it challenging to harmonise existing fields into a single estimate for the cortex(50). Confident that we can derive accurate estimates of cellular proportions in the adult cortex, we used our novel reference panels to characterise the cellular composition of the adult cortex using both datasets. In order to make inferences about the relative proportions of different subtypes of neurons and glial cells, we limited these comparisons to the estimates derived from reference panel 8, which contained the most specific combination of cell fractions using the IDOL method to select cell-specific sites. Plotting the distribution of cellular composition, we observe high levels of inter-individual variation (**Figure 5**, **Table 2**) across the samples. Glial cells were more abundant than neuronal cells (Exeter: mean neuronal proportion 0.336 (SD = 0.0627) vs mean glial proportion 0.683 (SD = 0.0723), Jaffe: mean neuronal proportion 0.309 (SD = 0.0582) vs mean glial proportion 0.711 (SD = 0.0692)). Within the neuronal cells, NeuNPos/SOX6Neg were more abundant on average (Exeter: mean = 0.306 (SD = 0.0557), Jaffe: mean = 0.295 (SD = 0.0555)) than NeuNPos/SOX6Pos cells (Exeter: mean = 0.0303 (SD = 0.0155), Jaffe: mean = 0.0132 (SD = 0.00989)). Within the glial cells, the NeuNNeg/SOX10Pos were most abundant on average (Exeter: mean proportion = 0.273 (SD = 0.153), Jaffe: mean proportion = 0.305 (SD = 0.133)) followed by the NeuNNeg/SOX10Neg/IRF8Neg (Exeter: mean proportion = 0.241 (SD = 0.0928), Jaffe: mean proportion = 0.232 (SD = 0.0807)). NeuNNeg/SOX10Neg/IRF8Pos was the least abundant predicted fraction ((Exeter: mean proportion = 0.169 (SD = 0.0388), Jaffe: mean proportion = 0.175 (SD = 0.0289)). The broad consistency across datasets in these relative abundance estimates supports the notion of an average pre-determined ratio of brain cells to underpin brain function but that this is highly variable across individuals. It is therefore, important to quantify cellular composition accurately for the purposes of controlling for potential confounding and may indeed be an interesting phenotype themselves in the study of brain development and brain disease.

**Figure 5.**
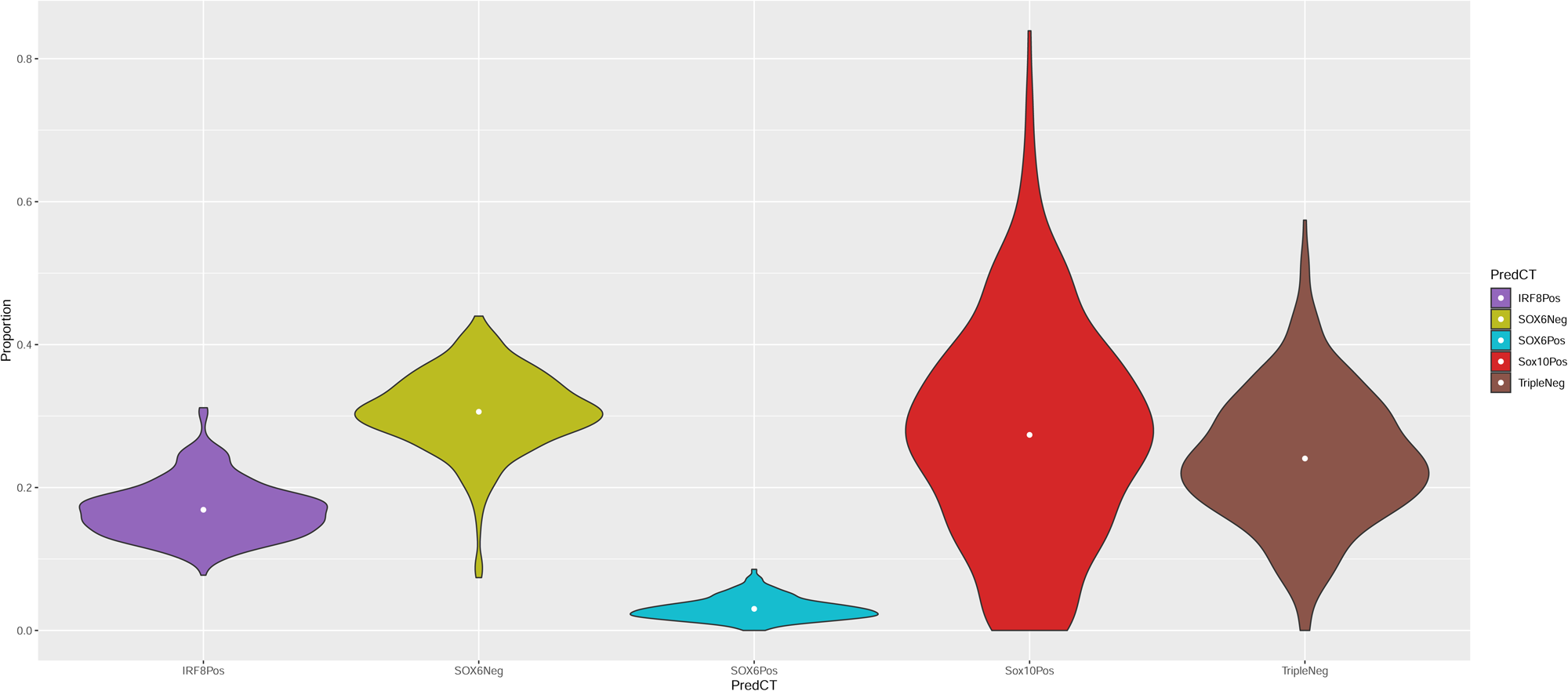
Cellular composition of adult prefrontal cortex. Violin plots of the distribution of proportion of brain cell types in the adult prefrontal cortex (n = 377), estimated using reference panel 8 and the IDOL algorithm for selecting cell-specific sites.

**Table 2.** Summary of proportions of cell types in the adult prefrontal cortex

| Cell types |  |  | Exeter |  | Jaffe |  |
| --- | --- | --- | --- | --- | --- | --- |
|  |  |  | Mean | SD | Mean | SD |
| Neuronal | all |  | 0.336 | 0.0628 | 0.309 | 0.0582 |
|  | NeuNPos/SOX6Neg | Excitatory (glutamatergic) neuronal enriched | 0.306 | 0.0557 | 0.295 | 0.0555 |
|  | NeuNPos/SOX6Pos | Inhibitory (gabaergic) neuronal enriched | 0.0303 | 0.0155 | 0.013 | 0.0099 |
| Glial | all |  | 0.683 | 0.0723 | 0.711 | 0.0692 |
|  | NeuNNeg/SOX10Neg/IRF8Pos | microglia enriched | 0.169 | 0.0388 | 0.175 | 0.0289 |
|  | NeuNNeg/SOX10Pos | oligodendrocyte-enriched | 0.274 | 0.153 | 0.305 | 0.133 |
|  | NeuNNeg/SOX10Neg/IRF8Neg | astrocyte enriched | 0.241 | 0.0928 | 0.232 | 0.0807 |

Exploring this further, were interested if there were any biological factors associated with the variation in cellular composition we observed. To streamline these analyses, we selected the optimal reference models for estimating the composition of each cell fraction (**Supplementary Table 8**), noting that correlations between fractions across panels were very high (**Supplementary Figure 20**). Testing the proportion of each cell type against age and sex, the only association that survived multiple testing in both datasets (P < 0.005, corrected for 10 cell types), was a higher proportion of NeuNPos/SOX6Pos cells in males (Exeter mean difference in males = 0.00229, P = 5.84×10^-5^; Jaffe mean difference in males = 0.00653, P = 6.97×10^-5^)(**Supplementary Table 9; Supplementary Figures 21-24**).

### The degree of Alzheimer’s disease neuropathology is associated with DNAm derived estimates of neuronal and glial composition

Finally, we were interested in whether the added specificity of our cellular composition estimates could enhance our understanding of the neuropathology of Alzheimer’s disease using data from recent analyses of DNAm differences associated with tau and amyloid pathology using bulk cortex(30). We estimated cellular proportions for each of the 10 fractions across three datasets where DNAm had been profiled in bulk prefrontal cortex tissue samples (total N = 864; **Supplementary Table 10**)(30, 42, 43). To ensure our subsequent analysis of cellular proportions were not biased, we first tested whether increasing tau pathology (quantified by Braak stage) influences the accuracy of cellular deconvolution. Although all models showed the same trend of decreasing CETYGO scores associated with increasing neurofibrillary tau tangles (**Supplementary Table 11**), only the CETYGO scores from reference panel 3 (mean change per Braak stage = -9.14×10^-4^,P = 8.24×10^-5^) were significantly related to pathology (P < 0.00333, corrected for 15 models). We found a significant association (P < 0.005, corrected for 10 cell types) for the prevalence of two estimated cell fractions with increasing levels of AD pathology (**Figure 6, Supplementary Table 12**). These data detected a decrease in the proportion of NeuNNeg/SO×10Neg nuclei (mean change per Braak stage = -0.00459, P = 0.00172), and an increase in the proportion of NeuNNeg/SO×10Pos nuclei (mean change per Braak stage 0.0744, P = 0.000555) with increasing tau pathology. There were also trends for significant negative correlations between the proportions of NeuNPos nuclei (mean change per Braak stage = -0.00282, P = 0.00993), SATB2Pos nuclei (mean change per Braak stage = - 0.00365, P = 0.00574) and NeuNPos/SOX6Pos (mean change per Braak stage = -0.00106, P = 0.00755) and a trend for a positive correlation with NeuNNeg (mean change per Braak stage = 0.0036, P = 0.006543).

**Figure 6.**
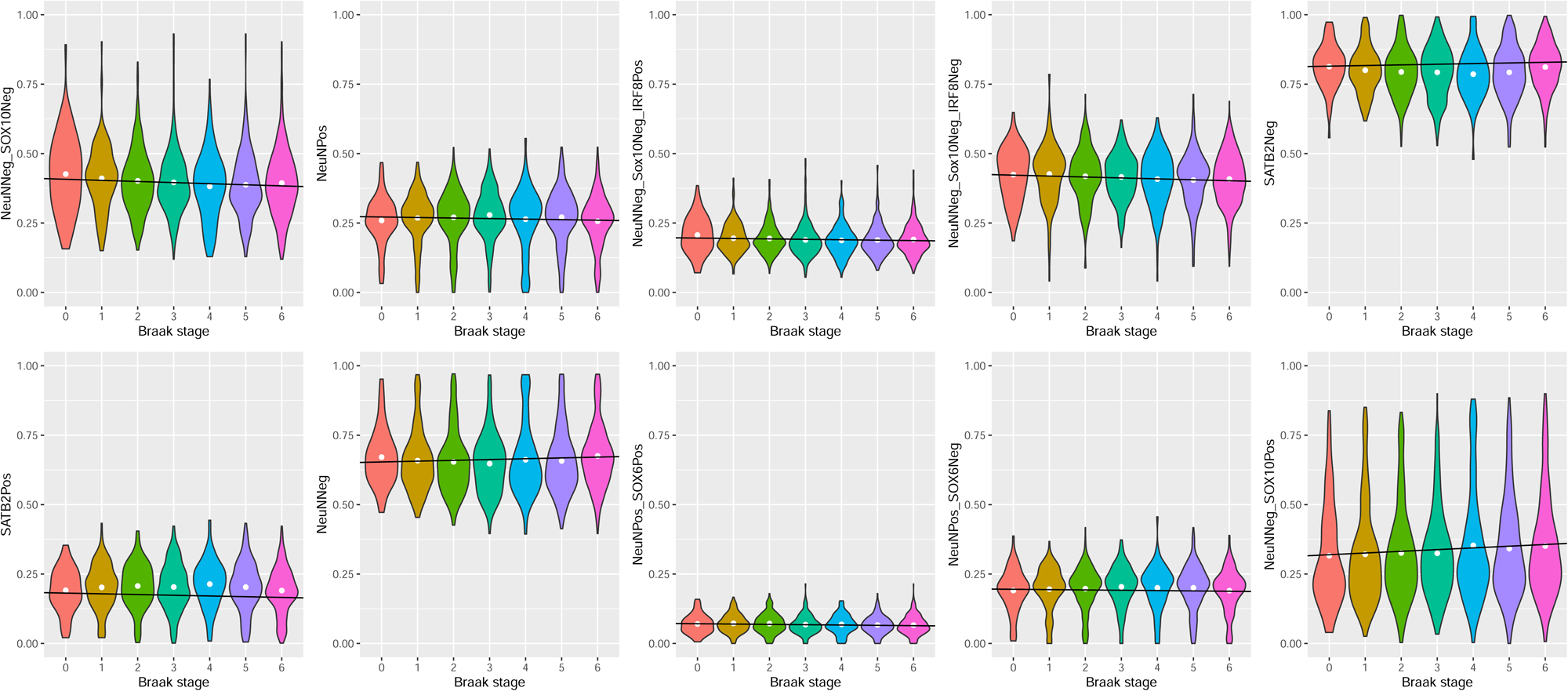
Cellular composition of adult prefrontal cortex varies as a function of Alzheimer’s Disease neuropathology. Violin plots of the distribution of proportion of brain cell types estimated from DNA methylation data generated from the prefrontal cortex (n = 864) grouped by Braak tangle stage. Each panel represents a different cell type, estimated using the optimal reference panel for that cell type (**Supplementary Table 8**).

## Discussion

We have generated genome-wide DNAm profiles for different cell types isolated from human cortex tissue, including novel profiles for several glial subtypes. We have demonstrated that these are applicable for use with established deconvolution algorithms and can be used to estimate cellular proportions in the cortex and other regions of the human brain from bulk DNAm data. Our proposed reference panel for deconvolution is the most extensive available for the human cortex and enables the prediction of neurons and glia, in addition to the prediction of two neuronal sub types (excitatory and inhibitory) and three glial subtypes (oligodendrocytes, microglia and astrocytes). We demonstrate that this approach produces accurate and informative estimates of cellular proportions from DNAm profiles generated using adult bulk human cortex samples but is not appropriate for the use in prenatal or cerebellum samples for which bespoke reference panels will be required.

Previous efforts to characterise the cellular composition of the human brain has produced a wide range of estimates, especially where the ratio of different cell types is concerned. This is in part due to the use of different methodologies, but perhaps more critically, due to the study of different brain regions and variation in whether the assay was limited to just the grey matter, white matter, or both(50). Our data could prove valuable in synthesising the existing research into a coherent conclusion. We observed approximately twice as many glial cells relative to neuronal cells, in line with the previously reported glial to neuronal ratio for cortical tissue consisting of both white and grey matter(50). Of neuronal cells we found that the proportion of GABAergic (inhibitory) neurons in the order of 5-10%, a bit lower compared to published literature stating this is between 10 and 20%(51). Within non-neuronal cells, we found that oligodendrocytes were the most frequent glial subtype, representing ∼40% of glial cells, followed by astrocytes (∼35%) and then microglia (∼25%). The rank ordering of abundance of glial subtypes is broadly consistent with the existing literature, although the estimated proportions differ, with a lower than expected proportion of oligodendrocytes and higher than expected proportion of microglia. We should caveat that our analysis of computationally constructed bulk profiles highlighted that the estimation of microglia proportion is better than the estimation of oligodendrocyte proportion and the estimation of astrocyte proportion is worst. Critically, our data highlight large variation in the composition of different cell types across samples, consistent with previous deconvolution studies of brain(16) and studies of cellular heterogeneity using other methods(50), reinforcing the importance of including these variables as covariates in association analyses(9).

As well as being potential confounders, there is interest in using these variables as phenotypes in epidemiological studies to identify the sources of the variation. We tested for effects of age and sex, and only found nominal associations between sex and one cell type, inhibitory neurons. To establish the biological validity of these cellular composition variables we tested them against semi-quantitative measures of Alzheimer’s disease neuropathology in existing datasets generated by our group(30, 42, 45). Our data was consistent with the known neuropathological effects of neuronal loss observed with the progression of Alzheimer’s disease(10, 11, 52), highlighting a decrease in the proportion of neurons observed in both inhibitory and excitatory neurons. We also detected changes in the composition of glial cells with the proportion of oligodendrocytes increasing and the proportions of microglia and astrocytes decreasing as tau tangles accumulate in the brain. This finding does not contradict reports that astrocytes and microglia exhibit enhanced activity in Alzheimer’s disease(12, 13, 53). Cellular deconvolution harnesses sites in the genome where there are cell-specific DNAm signatures that define cell identity (i.e. ubiquitous across all cells of that type) and likely does not capture changes in activation state (which potentially varies across a population of cells). One of the limitations of the methodology is it only allows us to determine cellular proportions rather than abundances. Given that the proportion of one cell type is influenced by the abundance of all cell types, significant associations with the proportion of an individual cell type might not be due to changes in the abundance of that cell type but changes in the overall composition. For this reason, caution needs to be applied when interpreting significant associations with these variables.

Given the use of four different FANS gating strategies to obtain different populations of nuclei, we had reference data for 10 different fractions of brain cell types, where some of these fractions targeted overlapping sets of nuclei. For this reason, we were able to propose 8 different ways to combine these data into reference panels for cellular deconvolution, with 6 of these reference panels consisting of non-overlapping fractions of nuclei. This is therefore, the most comprehensive study to date investigating how the composition of different reference panels affects the estimation of cellular heterogeneity. While our novel reference panel is primarily of interest to those studying variable DNAm in brain disorders, our analyses provide broader insights into the methodology of cellular deconvolution that are application for studies involving any bulk tissue.

It is reasonable to assume that the optimal reference panel would have the most diverse and specific set of cell types available, and our data demonstrate subtle improvements in accuracy when using models that contain a more specific set of subtypes. In addition to comparing different reference panels we also compared two methods for selecting cell specific sites (i.e. how the deconvolution model itself is trained) using an ANOVA or the IDOL algorithm(19), although this did not introduce much variation in performance. We found larger differences in performance between cell types and between reference panels than between training methodologies. We conjecture that this is due to variation in the quality of the reference data for each cell type, which is affected by both the signal-to-noise ratio of the DNAm array data and the efficiency of the isolation of those cell types. We were able to classify the different fractions into three performance tiers. The top tier with near perfect performance in our simulations included NeuNPos (neuronal enriched), NeuNPos/SOX6Pos (GABAergic neuronal enriched), NeuNPos/SOX6Neg (glutamatergic neuronal enriched) and NeuNNeg (glial enriched). The next tier, also associated with high accuracy statistics, included NeuNNeg/SO×10Neg (microglial & astrocyte enriched), NeuNNeg/SO×10Neg/IRF8Pos (microglial enriched), SATB2Pos (excitatory neuronal enriched), and SATB2Neg (inhibitory neuronal and glial enriched). The third tier included NeuNNeg/SO×10Pos (oligodendrocyte enriched) and NeuNNeg/SO×10Neg/IRF8Neg (astrocyte enriched) which were associated with a noticeable drop in performance metrics. While they likely still function as valuable proxies for variation in composition associated with these cell types, they are potentially affected by more noise, which will negatively affect the power to detect between-sample differences with these cell types. We expected that positively selected fractions (i.e. where an antibody is used to isolate a subset of nuclei) would be associated with higher degree of accuracy than negatively selected fractions (i.e. the population of unstained nuclei) due to increased levels of heterogeneity. This was not always the case, with the NeuNNeg/SO×10Neg fraction predicted more accurately than NeuNNeg/SO×10Pos fraction. Even within a purified population of nuclei, there is likely to be a heterogeneous mixture of different cellular subtypes and the extent of this heterogeneity will vary depending on the class of cell types and the activation state of any given cell. Another factor influencing the accuracy of the estimates of particular cell types is the availability of DNAm sites in the dataset that differentiate cell types. As has been shown for cell types in whole blood(54), our data confirmed that the magnitude of differences between brain cells is largely a function of their lineage. In other words, the major source of variation in these data was captured differences between the two major classes of brain cells, neurons and glial. The subsequent lower order sources of variation then captured the differences within these classes (e.g. astrocytes from oligodendrocytes). Interestingly, microglia, which arise from an entirely different lineage compared to the other brain cell types, sit within the glial cluster. There are fewer (and smaller) differences between more developmentally-related cell types to harness for deconvolution, making the analysis more difficult. This highlights a potential limitation of using microarray technology; having genuinely genome-wide DNAm data would likely be an advantage for or even essential for further resolving the cellular heterogeneity of the brain further into more specialised cell types.

When characterising the performance of estimates of cellular composition there are two statistical properties to consider. First is the absolute accuracy, which is important if the objective is to make inferences about the cellular profile of the brain. Second is the ability to capture a gradient of variation, i.e. the correlation. This is important if the aim is to test for associations with other phenotypes or use as covariates in analyses. When deciding which set of cellular composition variables to use, it is worth considering what they are going to be used for. If the objective is to test for associations between each cell type and an outcome (or adjusting for this variation) then it would be logical to select the most accurate estimate for each cell type, even if this means using different models for different cell types. The consequence of this approach is that the sum across all the cell types will not total 1.

When comparing the performance of different reference panels we have demonstrated how our accuracy metric for cellular deconvolution, CETYGO(55), can be applied. Our results reinforce the conclusions from the original work, that the parameters of the distribution of the CETYGO score are reference panel and technology specific. The association in the analyses between batch and accuracy highlight that data quality are important not only for increasing power to detect significant effects with an outcome, but also to effectively capture cellular heterogeneity. We therefore, recommend that not only do future studies take advantage of our expanded set of brain cell type composition variables, but that they also include the CETYGO score as part of their quality control to identify outlier samples.

## Conclusions

In summary, we have generated an expanded set of reference data for the purpose of estimating the cellular heterogeneity of DNAm profiles generated from bulk human cortex tissue. These variables will be critical covariates to include in future epigenetic studies of brain disorders to minimise the risk of false positive associations and improve our understanding of the changes in the brain that underpin the development of psychiatric disorders and neurodegenerative diseases.

## Methods

### Isolation of neural nuclei from post-mortem brain tissue

Post-mortem prefrontal cortex (PFC) samples were processed using our optimised FANS protocol(31). PFC post-mortem brain tissue from 43 adult donors (aged 55-95 years old) was provided from 9 brain banks from the UK, Canada and US (Brains for Dementia Research Network of Brain Banks, King’s College London, Harvard, UCLA, Oxford, Miami, Douglas Bell, Pittsburgh and Mount Sinai Brain Banks). Human cortex tissue was collected under approved ethical regulation at each centre and transferred to our care through Materials Transfer Agreements.

500 mg of frozen brain tissue was homogenised in lysis buffer (2mL) using a pre-chilled Dounce homogeniser. The homogenate was layered above 8mL of sucrose solution in ultracentrifuge tubes (Thermo Scientific, Cat N# 03699) (1mL per tube) and overlaid with a lysis buffer (2 mL per tube) for a final volume of 11 mL. Following the purification of nuclei by density gradient ultracentrifugation (model: Sorvall™ WX 80+; rotor: TH-641; speed: 108670.8 x g, 45 min at 4°C) each nuclei pellet was resuspended in 1 mL staining buffer and incubated on ice for 10 min. Nuclei suspensions were then pelleted in 2mL Eppendorf tube (DNase, RNase free) by centrifugation at 1000xg, 5 min at 4°C. After carefully discarding the supernatant, nuclei pellets were resuspended in fresh staining buffer and pooled together. After adding 2 µL of DNA dye (Hoechst 33342, Abcam, Cat # ab228551) 150μL of nuclei solution (Hoechst only), was transferred in a new 2mL tube and volume made up to 1mL with fresh SB for use as the Unstained Control. For the “Stained” tube, the volume removed was replaced with fresh staining buffer and the suspension was then immune-stained with a combination of antibodies including, NeuN-Alexa488, anti-SO×10 NL577-conjugated and/or anti-IRF8 APC-conjugated antibodies. Details of the three different gating strategies we implemented can be seen in **Supplementary Figure 1**. Both stained and unstained tubes were incubated for 1.5 hours on a spinning rotor in the dark at 4 °C. Tubes were spun at 1000 x g, for 5min at 4°C and the supernatant was carefully discarded from both tubes and remaining nuclei pellets were re-suspend in staining buffer (500µl unstained, 1 - 1.5mL stained tube - dependent on pellet density) using wide bore tips. Tubes were brought to the FACS Aria III cell sorter and kept on ice for the entire procedure of machine setup and sorting. Nuclei suspensions were assessed for the presence of debris by adjusting the gating strategy appropriately before proceeding with nuclei capture. The 100 µM nozzle was used and the event rate during data acquisition and sample collection was kept ≤ 3000 events/sec. On average, for each sorted population, 200,000 nuclei were collected for extraction of genomic DNA.

### DNA extraction

Nuclei aliquots were defrosted on ice and 50 mL the volume of the aliquots made up to 1mL with Slagboom buffer (SB) (5mL 10x STE buffer, 5mL 5% SDS, 40mL RNase-free DNase-free water)). Nuclei were collected and stored in FACSFlow buffer, approximately 600µL per tube for 200,000 nuclei. 1μL of DNase free RNase-A (10 mg/ml) per 500μL of sample was added and the samples were incubated at 37°C for 45 min (heat block). 5μL of proteinase K (20 mg/ml) (Thermo Fisher Scientific, Waltham, MA, USA) was then added and the samples were inverted at least 10 times. The samples were then incubated at 60°C for 1 hour, and then cooled to room temperature (RT) for 5 min. 200μL of “Majik Mix” (a proprietary reagent made from 1:1 ratio yeast Reagent 3 (Autogen Bioclear, Caine, Wiltshire, UK) and 100% ethanol) was added and the samples were mixed by vigorous inversions before being centrifuged at 17,000xg for 10 minutes at RT. For each sample the supernatant was carefully recovered and transferred to a new labelled tube (50μL was left at the bottom of each tube). Another 200μL of Majiik Mix were added to each tube and samples were again mixed by vigorous inversion before being centrifuged at 17,000xg for 10 minutes at RT. The upper layer of each tube was carefully recovered (making sure to leave approximately 50μL to prevent carrying over any of the lower layer) and transferred to a new appropriately labelled tube. Where exceeding 1mL total volume, supernatant was equally distributed into 2 new tubes. An equal volume of 100% Isopropanol (Sigma-Aldrich Corporation, St. Louis, MO, USA) was added to each sample (e.g. 1mL supernatant + 1mL 100% Isopropanol) and slowly mixed by inverting to precipitate the DNA. At this stage, 0.5-0.8μL GlycoBlue™ Co-precipitant (Invitrogen Ltd, Inchinnan, UK) was added to each sample. When a typical acetate/alcohol precipitation is done, the GlycoBlue™ Coprecipitant will precipitate with the nucleic acids, facilitating good DNA recovery while increasing the size and visibility of the pellet. The samples were then mixed by inverting the tubes ∼10 times and centrifuged at 17,000xg for 15 minutes at RT. For each tube the supernatant was carefully removed and discarded. 500µL of 80% ethanol were added to each tube, samples were then mixed gently and centrifuged at 17,000xg for 5 min. The supernatant was carefully removed and the pellets were left to air dry for 20 minutes or until dry. Each DNA pellet was resuspended in 15µL of RNAse, DNase-free water and left at 4°C overnight to fully dissolve before quantification.

### Methylomic profiling

500ng of genomic DNA from each sample was treated with sodium BS using the Zymo EZ-96 DNA Methylation-Gold™ Kit (Cambridge Bioscience, UK) according to the manufacturer’s standard protocol. All samples were then processed using the EPIC 850K array (Illumina Inc, CA, USA) according to the manufacturer’s instructions, with minor amendments and quantified using an Illumina HiScan System (Illumina, CA, USA). Individuals were randomised and sorted fractions from the same individual and FACs gating run were processed on the same BeadChip, where within a BeadChip the location of each fraction was randomised. In total 42 NeuNPos, 39 NeuNNeg/SO×10Pos, 33 NeuNNeg/SO×10Neg, 12 SATB2Pos, 19 NeuNNeg/SO×10Neg/IRF8Neg, 34 NeuNNeg/SO×10Neg/IRF8Pos, and 9 SATB2Neg samples were run on the DNAm arrays.

### DNAm data preprocessing

DNA methylation data was loaded in R (version 3.6.3) from idat files using the package bigmelon(56). These data were processed through a standard quality control pipeline which included the following steps: 1) checking methylated and unmethylated signal intensities, excluding samples where this was < 500; 2) using the control probes to ensure the sodium bisulfite conversion was successful, excluding any samples with median < 80; 3) use of the 59 SNP probes to confirm that samples from the sample individual were genetically identical; 4) pfilter function from wateRmelon package to exclude samples with > 1% of probes with detection P-value > 0.05 and probes with > 1% of samples with detection P-value > 0.05 5) counting the number of missing values per sample and excluding samples with > 2% probes missing.

To confirm the success of the FANS sorting we applied a bespoke classification algorithm based on principal components analysis across all autosomal DNA methylation sites. The general objective was to compare each sample to the average profile of the labelled sample type. We Studentized the values of the first two principal components and excluded those above a threshold of 1.5, to minimise the effects of outliers (which are likely to be due to either mislabelling or suboptimal FANS sorting) on the average profiles for each cell type. For each cell type, we then calculated the mean and SD of the first two principal components only including the non-outlier samples. These were then used to calculate sample level scores that captured the similarity of the observed sample and the expected profile for that cell type. This was defined as the value of the principal component for that sample minus the mean for the cell type divided by the standard deviation for the cell type. The value can be interpreted as the number of SD from the mean that sample is, where lower values are desirable. This was performed separately for the first two principal components and then combined into a single score by taking the maximum, referred to here as the maxSD score. Prior to confirming the labelling of individual samples, we first wanted to confirm at an individual level that we had successfully isolated distinct fractions of nuclei. For this we calculated individual level metrics that represents the efficiency of the FANS sort. These are defined as the median across all the maxSD scores for that individual. Where the FANS sorting worked well (i.e. all antibodies were stained and gated accurately), all the samples from that individual should be close to their relevant average profile and this score will be low. Where the FANS sorting for an individual did not successfully isolate the relevant cell types, these samples will still be heterogeneous mixtures of cells, and sit in the middle of the principal component space, far away from their average profile, all with large maxSD. By taking the median, we ensure that we focus here on detecting FANS sorts, where the separation into specific cell types was not successful, rather than instances where just one/two samples are affected/mislabelled. Visual inspection of the best and worst performing individuals, informed us that a threshold of 5 was appropriate. It is also important to exclude these individuals prior to performing the cell type checking at a sample level as it enables us to ensure we have high signal to noise average profiles for the cell types. Having excluded all the samples associated with any individual deemed to have inefficient FANS sorting, we recalculated the Studentized values prior to recalculating the cell type means and SD for the first two principal components. Samples were retained if their principal components values were within two standard deviations of the mean of their labelled cell type. Samples that were more than two standard deviations away from the mean in either of the first two principal components were excluded from further analyses. Samples were then normalised using the dasen function(57), separately for each cell type.

### EPIGABA data

DNA methylation data generated with Illumina 450K BeadChip array were downloaded from the Synapse portal (syn7072866) for 5 NeuNPos/SOX6Pos, 5 NeuNPos/SOX6Neg, and 5 NeuNNeg samples. idat files for these samples were put through the same quality control pipeline described above, and the same classification algorithm was performed to confirm successful isolation and high quality reference data for the purpose of cellular deconvolution.

### Merging reference DNA methylation datasets

Our Exeter reference dataset was joined to the EpiGABA reference dataset. Given the use of two different technologies, we filtered to sites common to both the EPICarray and 450K array. Prior to training any deconvolution models, these datasets were filtered to only include autosomal DNAm sites and remove cross-hybridising probes and SNP probes as defined in publicly available resources(58, 59).

### Generation of deconvolution models and selection of cell-specific sites

Given the range of different cell types we have isolated, and the fact that these represent overlapping sets of nuclei, we defined 8 different combinations of cell types each of which represent a different reference panel (**Supplementary Figure 3**). To test and compare the performance of these panels against a known truth we trained a series of Houseman constraint projection deconvolution models using our novel reference data. These were then tested against reconstructed brain tissue DNAm profiles where we combined cell-specific profiles in a weighted linear sum of pre-specified proportions of each cell type. For each simulation, one sample for each cell type was removed to generate the testing data, and the remaining samples formed the training data, such that the train and test data consisted of distinct sets of samples. It should be noted though, that they were from the same experimental batch, and plausibly share technical, batch-specific effects. In this framework, training the models essentially means selecting the cell-specific sites that form the basis of the deconvolution algorithm. We used two different methods to select these sites. First, an ANOVA is performed across all samples in the training data to identify sites that are significantly different (p value < 1×10^-8^) between the brain cell types, selecting 2N sites per cell type (N hypermethylated and N hypomethylated). This is the approach implemented by minfi (via the *EstimateCellCounts* function)(60). The second approach, is the IDOL method(19). This also starts with an ANOVA to identify a larger pool of possible cell-specific sites, in our case the default selection of 150 sites per cell type with smallest and largest t-statistics. It then tests random subsets of sites to refine this list to a smaller set of size M probes such that the optimal performance is achieved. To determine whether a particular subset of sites is a better fit than the current best subset, it requires a separate set of reconstructed test profiles with known cellular composition. These were constructed as described below for our testing data but from a sample selected at random from the training samples. The effect of individual CpGs on the accuracy of the deconvolution is assessed by comparing the accuracy of estimated cellular composition with and without that CpG. CpGs that confer a positive effect are then up weighted such that they are more likely to be selected in the next random subset. The selection of random subset of sites was performed a maximum of 300 times. This optimisation was performed using the *IDOLoptimize()* function provided in the IDOL R package. Note the IDOL method is only applicable to reference panels with > 2 cell types. Therefore from our 8 reference panels there were 15 trained models. For each of these models we trained the models multiple times to select between 20 and 200 probes per cell type, increasing in units of 20 probes.

### Generation simulated bulk brain profiles

To construct bulk brain profiles for testing, we combined the cell specific test profiles in fixed proportions that represented the full spectrum of possible combinations. Each reference panel was only tested against reconstructed profiles consisting of the same cell types. Cell type proportions were increased in 0.1 units, where each cell type represented at least 0.1, up to a maximum of 0.9 and such that the total of all cell type proportions equalled 1. As each reference panel consists of different numbers of cell types, the possible number of reconstructed profiles tested differs by virtue of the different number of combinations possible with that number of cell types. DNAm levels in the test data at these cell-specific sites are then computed into estimates of cellular proportions using a quadratic programming methodology as described by Houseman(15). This process was repeated for 10 different train-test splits of the reference data. This methodology was implemented using functions in the CETYGO package(55) which are adaptations of functions from the minfi package(60) that takes matrices of beta values as input for the training and testing data.

### Training deconvolution models for use with empirical bulk brain profiles

To train the deconvolution algorithm for all 15 models for use with empirical bulk brain datasets, and for sharing with the wider research community, we used all available samples for each cell type (**Supplementary Table 1**). Cell-specific sites were selected in the same way as above with a total of 100 probes per cell type selected (i.e. for a two cell type reference panel up to 200 probes were selected).

### Profiling the performance of neural cell type deconvolution in empirical datasets

We used four datasets of bulk brain DNAm profiles from two sources to further characterise the performance of the neural reference panels. The first source contains data generated by our group at the University of Exeter (www.epigenomicslab.com) across a range of projects and includes i) a dataset of 377 adult PFC samples (BA9)(40–43), ii) a dataset of 851 adult samples from 9 other brain regions including additional cortical regions, the striatum, the hippocampus, and cerebellum (**Supplementary Table 6**)(39–43, 46), and 167 prenatal and childhood samples(39, 48). The second source is the publicly available data provided by Jaffe et al(44) and includes 415 adult PFC samples. All datasets were processed by our group through a standard QC pipeline(30) and were normalised using the *dasen* function in the *wateRmelon* package(61). Cellular composition was estimated for all samples using all 15 models and then selecting the estimates from the best performing models for each cell type (**Supplementary Table 8**).

In the adult PFC datasets we used a linear regression model to test for batch effects (slide) and biological (age, age^2^ and sex) effects on the CETYGO score. P-values for the age, age^2^ and sex covariates were taken from t-tests of the estimated regression coefficients. The P-value for the batch effect was taken from an ANOVA comparing with full model to a nested model without the batch covariate. In the Exeter adult multi-tissue dataset we tested for brain region effects on the CETYGO scores using a linear regression model, where PFC(BA9) was set to the baseline, so that we estimated coefficients and p-values for all other brain regions relative to the PFC. In this analysis we controlled for age, age^2^, sex but not batch as data generated from different brain regions were run in different batches. To test for effects on cellular composition we used the same linear regression models as described here for the estimated proportion of each cell type in turn.

### Testing the associations between cellular composition and Alzheimer’s disease neuropathology

We additionally generated estimates of cellular composition in three in house Alzheimer’s disease DNA methylation datasets(30, 42, 43), where data had been generated from DNA extracted from the PFC (**Supplementary Table 10**). Cellular composition was estimated for all samples using all 15 models and then selecting the estimates from the best performing models (**Supplementary Table 8**). We used a linear regression model within each cohort to test for associations between Braak stage (modelled as a continuous variable) and either the CETYGO score or estimated proportion of each cell type, including covariates for age and sex. The estimated coefficients for Braak stage and the associated standard errors were then meta-analysed together using the R package meta(62). Given that we only included three studies, we present only the fixed effect results in the main text, but the random effect results are also available in the relevant Supplementary Tables.

## Declarations

### Ethics approval and consent to participate

Ethical approval for the study was granted by the University of Exeter Medical School Research Ethics Committee (13/02/009).

### Consent for publication

Not applicable

### Availability of data and materials

Data generated for this project are available at NCBI Gene Express Omnibus (GEO) under accession number GSE234520. We also reanalysed data previously made available via GEO (via accession numbers GSE74193, GSE59685, GSE80970, GSE88890, GSE43414) and the synapse platform (syn7072866, syn8263588). Code for the quality control of the DNAm data can be found on GitHub at https://github.com/ejh243/BrainFANS/tree/master/array/DNAm/preprocessing. The code for analyses presented here can be found on GitHub at https://github.com/ejh243/BrainFANS/tree/master/array/DNAm/analysis/neuralCellComposition. Our new trained deconvolution models for brain are made available to the wider research community via our R package CETYGO available on GitHub (https://github.com/ds420/CETYGO).

### Competing interests

The authors declare that they have no competing interests.

### Funding

These data were generated as part of Medical Research Council grant K013807 to JM and Alzheimer’s Research UK (ARUK) grant ARUK-PPG2018A-010 to E.D. E.H., J.M., E.L.D, and L.C.S were supported by Medical Research Council (MRC) grants K013807 and W004984 (awarded to J.M.). E.H is supported by an Engineering and Physical Sciences Research Council Fellowship EP/V052527/1. Data analysis was undertaken using high-performance computing supported by a Medical Research Council (MRC) Clinical Infrastructure award (M008924) to J.M. The EpiGABA DNA methylation data were generated as part of the PsychENCODE Consortium, supported by: U01DA048279, U01MH103339, U01MH103340, U01MH103346, U01MH103365, U01MH103392, U01MH116438, U01MH116441, U01MH116442, U01MH116488, U01MH116489, U01MH116492, U01MH122590, U01MH122591, U01MH122592, U01MH122849, U01MH122678, U01MH122681, U01MH116487, U01MH122509, R01MH094714, R01MH105472, R01MH105898, R01MH109677, R01MH109715, R01MH110905, R01MH110920, R01MH110921, R01MH110926, R01MH110927, R01MH110928, R01MH111721, R01MH117291, R01MH117292, R01MH117293, R21MH102791, R21MH103877, R21MH105853, R21MH105881, R21MH109956, R56MH114899, R56MH114901, R56MH114911, R01MH125516, and P50MH106934 awarded to: Alexej Abyzov, Nadav Ahituv, Schahram Akbarian, Alexander Arguello, Lora Bingaman, Kristin Brennand, Andrew Chess, Gregory Cooper, Gregory Crawford, Stella Dracheva, Peggy Farnham, Mark Gerstein, Daniel Geschwind, Fernando Goes, Vahram Haroutunian, Thomas M. Hyde, Andrew Jaffe, Peng Jin, Manolis Kellis, Joel Kleinman, James A. Knowles, Arnold Kriegstein, Chunyu Liu, Keri Martinowich, Eran Mukamel, Richard Myers, Charles Nemeroff, Mette Peters, Dalila Pinto, Katherine Pollard, Kerry Ressler, Panos Roussos, Stephan Sanders, Nenad Sestan, Pamela Sklar, Nick Sokol, Matthew State, Jason Stein, Patrick Sullivan, Flora Vaccarino, Stephen Warren, Daniel Weinberger, Sherman Weissman, Zhiping Weng, Kevin White, A. Jeremy Willsey, Hyejung Won, and Peter Zandi.

### Authors’ contributions

EH, ELD, LCS & JM designed the study; JB, BC, JPD, GC, RB & SP generated the data; EH, EMW and AF analysed the data; EH & JM interpreted the data and drafted the manuscript. All authors reviewed the manuscript.

## Acknowledgements

We acknowledge the supply of samples from; The Cambridge Brain Bank, covered by current REC approval NRES 10/HO308/56; the Quebec Suicide Brain Bank at Douglas Mental Health University Institute, Canada; the MRC funded University of Edinburgh Brain & Tissue Bank; the Harvard Brain Tissue Resource Centre, which is supported by HHSN-271-2013-00030C; Brains for Dementia Research has ethics approval from London – City and East NRES committee 08/H0704/128+5; Brain Endowment Bank, at Miller School of Medicine, University of Miami; The Mount Sinai NBTR (NIH Brain and Tissue Repository), JJ Peters VA Medical Center; the Oxford Brain Bank, supported by the Medical Research Council (MRC), the NIHR Oxford Biomedical Research Centre and the Brains for Dementia Research programme, jointly funded by Alzheimer’s Research UK and Alzheimer’s Society; The Neuropathology Brain Bank at the University of Pittsburgh, School of Medicine Department of Psychiatry; The Stanley Medical Research Institute Brain Collection courtesy of Drs. Michael B. Knable, E. Fuller Torrey, Maree J. Webster, and Robert H. Yolken; The Human Brain and Spinal Fluid Resource Center (HBSFRC) NIH Neurobiobank. This study was supported by the National Institute for Health and Care Research Exeter Biomedical Research Centre. The views expressed are those of the author(s) and not necessarily those of the NIHR or the Department of Health and Social Care.

## Notes

https://www.ncbi.nlm.nih.gov/geo/

https://www.synapse.org/

### Competing Interest Statement

The authors have declared no competing interest.

## References

1. Pidsley R, Zotenko E, Peters TJ, Lawrence MG, Risbridger GP, Molloy P, et al. Critical evaluation of the Illumina MethylationEPIC BeadChip microarray for whole-genome DNA methylation profiling. Genome Biol. 2016;17(1):208.

2. Sandoval J, Heyn H, Moran S, Serra-Musach J, Pujana MA, Bibikova M, et al. Validation of a DNA methylation microarray for 450,000 CpG sites in the human genome. Epigenetics. 2011;6(6):692–702.

3. Mansell G, Gorrie-Stone TJ, Bao Y, Kumari M, Schalkwyk LS, Mill J, et al. Guidance for DNA methylation studies: statistical insights from the Illumina EPIC array. BMC Genomics. 2019;20(1):366.

4. Hannon E, Lunnon K, Schalkwyk L, Mill J. Interindividual methylomic variation across blood, cortex, and cerebellum: implications for epigenetic studies of neurological and neuropsychiatric phenotypes. Epigenetics. 2015;10(11):1024–32.

5. Hannon E, Mansell G, Walker E, Nabais MF, Burrage J, Kepa A, et al. Assessing the co-variability of DNA methylation across peripheral cells and tissues: Implications for the interpretation of findings in epigenetic epidemiology. PLoS Genet. 2021;17(3):e1009443.

6. Salas LA, Zhang Z, Koestler DC, Butler RA, Hansen HM, Molinaro AM, et al. Enhanced cell deconvolution of peripheral blood using DNA methylation for high-resolution immune profiling. Nat Commun. 2022;13(1):761.

7. Roadmap Epigenomics Consortium, Kundaje A, Meuleman W, Ernst J, Bilenky M, Yen A, et al. Integrative analysis of 111 reference human epigenomes. Nature. 2015;518(7539):317–30.

8. Campagna MP, Xavier A, Lechner-Scott J, Maltby V, Scott RJ, Butzkueven H, et al. Epigenome-wide association studies: current knowledge, strategies and recommendations. Clin Epigenetics. 2021;13(1):214.

9. Jaffe AE, Irizarry RA. Accounting for cellular heterogeneity is critical in epigenome-wide association studies. Genome Biol. 2014;15(2):R31.

10. Crews L, Masliah E. Molecular mechanisms of neurodegeneration in Alzheimer’s disease. Hum Mol Genet. 2010;19(R1):R12–20.

11. West MJ, Coleman PD, Flood DG, Troncoso JC. Differences in the pattern of hippocampal neuronal loss in normal ageing and Alzheimer’s disease. Lancet. 1994;344(8925):769–72.

12. Tejera D, Heneka MT. Microglia in Alzheimer’s disease: the good, the bad and the ugly. Curr Alzheimer Res. 2016;13(4):370–80.

13. Malm TM, Jay TR, Landreth GE. The evolving biology of microglia in Alzheimer’s disease. Neurotherapeutics. 2015;12(1):81–93.

14. Accomando WP, Wiencke JK, Houseman EA, Nelson HH, Kelsey KT. Quantitative reconstruction of leukocyte subsets using DNA methylation. Genome Biol. 2014;15(3):R50.

15. Houseman EA, Accomando WP, Koestler DC, Christensen BC, Marsit CJ, Nelson HH, et al. DNA methylation arrays as surrogate measures of cell mixture distribution. BMC Bioinformatics. 2012;13:86.

16. Guintivano J, Aryee MJ, Kaminsky ZA. A cell epigenotype specific model for the correction of brain cellular heterogeneity bias and its application to age, brain region and major depression. Epigenetics. 2013;8(3):290–302.

17. Teschendorff AE, Breeze CE, Zheng SC, Beck S. A comparison of reference-based algorithms for correcting cell-type heterogeneity in Epigenome-Wide Association Studies. BMC Bioinformatics. 2017;18(1):105.

18. Bell-Glenn S, Thompson JA, Salas LA, Koestler DC. A Novel Framework for the Identification of Reference DNA Methylation Libraries for Reference-Based Deconvolution of Cellular Mixtures. Front Bioinform. 2022;2.

19. Koestler DC, Jones MJ, Usset J, Christensen BC, Butler RA, Kobor MS, et al. Improving cell mixture deconvolution by identifying optimal DNA methylation libraries (IDOL). BMC Bioinformatics. 2016;17:120.

20. Newman AM, Liu CL, Green MR, Gentles AJ, Feng W, Xu Y, et al. Robust enumeration of cell subsets from tissue expression profiles. Nat Methods. 2015;12(5):453–7.

21. Houseman EA, Molitor J, Marsit CJ. Reference-free cell mixture adjustments in analysis of DNA methylation data. Bioinformatics. 2014;30(10):1431–9.

22. Leek JT, Storey JD. Capturing heterogeneity in gene expression studies by surrogate variable analysis. Plos Genet. 2007;3(9):1724–35.

23. Rahmani E, Schweiger R, Rhead B, Criswell LA, Barcellos LF, Eskin E, et al. Cell-type-specific resolution epigenetics without the need for cell sorting or single-cell biology. Nat Commun. 2019;10(1):3417.

24. Zou J, Lippert C, Heckerman D, Aryee M, Listgarten J. Epigenome-wide association studies without the need for cell-type composition. Nat Methods. 2014;11(3):309–11.

25. Qi L, Teschendorff AE. Cell-type heterogeneity: Why we should adjust for it in epigenome and biomarker studies. Clin Epigenetics. 2022;14(1):31.

26. Lake BB, Ai R, Kaeser GE, Salathia NS, Yung YC, Liu R, et al. Neuronal subtypes and diversity revealed by single-nucleus RNA sequencing of the human brain. Science. 2016;352(6293):1586–90.

27. Herring CA, Simmons RK, Freytag S, Poppe D, Moffet JJD, Pflueger J, et al. Human prefrontal cortex gene regulatory dynamics from gestation to adulthood at single-cell resolution. Cell. 2022;185(23):4428–47.e28.

28. Nott A, Schlachetzki JCM, Fixsen BR, Glass CK. Nuclei isolation of multiple brain cell types for omics interrogation. Nat Protoc. 2021;16(3):1629–46.

29. Matevossian A, Akbarian S. Neuronal nuclei isolation from human postmortem brain tissue. J Vis Exp. 2008(20).

30. Shireby G, Dempster E, Policicchio S, Smith RG, Pishva E, Chioza B, et al. DNA methylation signatures of Alzheimer’s disease neuropathology in the cortex are primarily driven by variation in non-neuronal cell-types. bioRxiv. 2022:2022.03.15.484508.

31. Policicchio SSS, Davies JP, Chioza B, Jeffries A, Burrage J, Mill J, et al. DNA Extraction from FANS sorted nuclei. protocols.io 2020.

32. Kim J, Hannibal L, Gherasim C, Jacobsen DW, Banerjee R. A human vitamin B12 trafficking protein uses glutathione transferase activity for processing alkylcobalamins. J Biol Chem. 2009;284(48):33418–24.

33. Stolt CC, Rehberg S, Ader M, Lommes P, Riethmacher D, Schachner M, et al. Terminal differentiation of myelin-forming oligodendrocytes depends on the transcription factor So×10. Genes Dev. 2002;16(2):165–70.

34. Masuda T, Tsuda M, Yoshinaga R, Tozaki-Saitoh H, Ozato K, Tamura T, et al. IRF8 is a critical transcription factor for transforming microglia into a reactive phenotype. Cell Rep. 2012;1(4):334–40.

35. Huang Y, Song NN, Lan W, Hu L, Su CJ, Ding YQ, et al. Expression of transcription factor Satb2 in adult mouse brain. Anat Rec (Hoboken). 2013;296(3):452–61.

36. Akbarian S, Liu C, Knowles JA, Vaccarino FM, Farnham PJ, Crawford GE, et al. The PsychENCODE project. Nat Neurosci. 2015;18(12):1707–12.

37. Kozlenkov A, Wang M, Roussos P, Rudchenko S, Barbu M, Bibikova M, et al. Substantial DNA methylation differences between two major neuronal subtypes in human brain. Nucleic Acids Res. 2016;44(6):2593–612.

38. Seiler Vellame D, Castanho I, Dahir A, Mill J, Hannon E. Characterizing the properties of bisulfite sequencing data: maximizing power and sensitivity to identify between-group differences in DNA methylation. BMC Genomics. 2021;22(1):446.

39. Wong CCY, Smith RG, Hannon E, Ramaswami G, Parikshak NN, Assary E, et al. Genome-wide DNA methylation profiling identifies convergent molecular signatures associated with idiopathic and syndromic autism in post-mortem human brain tissue. Hum Mol Genet. 2019;28(13):2201–11.

40. Viana J, Hannon E, Dempster E, Pidsley R, Macdonald R, Knox O, et al. Schizophrenia-associated methylomic variation: molecular signatures of disease and polygenic risk burden across multiple brain regions. Hum Mol Genet. 2016.

41. Pidsley R, Viana J, Hannon E, Spiers HH, Troakes C, Al-Saraj S, et al. Methylomic profiling of human brain tissue supports a neurodevelopmental origin for schizophrenia. Genome Biol. 2014;15(10):483.

42. Lunnon K, Smith R, Hannon E, De Jager PL, Srivastava G, Volta M, et al. Methylomic profiling implicates cortical deregulation of ANK1 in Alzheimer’s disease. Nat Neurosci. 2014;17(9):1164–70.

43. Smith RG, Hannon E, De Jager PL, Chibnik L, Lott SJ, Condliffe D, et al. Elevated DNA methylation across a 48-kb region spanning the HOXA gene cluster is associated with Alzheimer’s disease neuropathology. Alzheimers Dement. 2018.

44. Jaffe AE, Gao Y, Deep-Soboslay A, Tao R, Hyde TM, Weinberger DR, et al. Mapping DNA methylation across development, genotype and schizophrenia in the human frontal cortex. Nat Neurosci. 2015.

45. Smith RG, Pishva E, Shireby G, Smith AR, Roubroeks JAY, Hannon E, et al. A meta-analysis of epigenome-wide association studies in Alzheimer’s disease highlights novel differentially methylated loci across cortex. Nat Commun. 2021;12(1):3517.

46. Murphy TM, Crawford B, Dempster EL, Hannon E, Burrage J, Turecki G, et al. Methylomic profiling of cortex samples from completed suicide cases implicates a role for PSORS1C3 in major depression and suicide. Transl Psychiatry. 2017;7(1):e989.

47. Jeffries AR, Mill J. Profiling Regulatory Variation in the Brain: Methods for Exploring the Neuronal Epigenome. Biol Psychiatry. 2017;81(2):90–1.

48. Spiers H, Hannon E, Schalkwyk LC, Smith R, Wong CC, O’Donovan MC, et al. Methylomic trajectories across human fetal brain development. Genome Res. 2015;25(3):338–52.

49. Alcamo EA, Chirivella L, Dautzenberg M, Dobreva G, Fariñas I, Grosschedl R, et al. Satb2 regulates callosal projection neuron identity in the developing cerebral cortex. Neuron. 2008;57(3):364–77.

50. von Bartheld CS, Bahney J, Herculano-Houzel S. The search for true numbers of neurons and glial cells in the human brain: A review of 150 years of cell counting. J Comp Neurol. 2016;524(18):3865–95.

51. Sahara S, Yanagawa Y, O’Leary DD, Stevens CF. The fraction of cortical GABAergic neurons is constant from near the start of cortical neurogenesis to adulthood. J Neurosci. 2012;32(14):4755–61.

52. Niikura T, Tajima H, Kita Y. Neuronal cell death in Alzheimer’s disease and a neuroprotective factor, humanin. Curr Neuropharmacol. 2006;4(2):139–47.

53. Heneka MT, Kummer MP, Latz E. Innate immune activation in neurodegenerative disease. Nat Rev Immunol. 2014;14(7):463–77.

54. Hannon E, Mansell G, Burrage J, Kepa A, Best-Lane J, Rose A, et al. Assessing the co-variability of DNA methylation across peripheral cells and tissues: implications for the interpretation of findings in epigenetic epidemiology. bioRxiv. 2020:2020.05.21.107730.

55. Vellame DS, Shireby G, MacCalman A, Dempster EL, Burrage J, Gorrie-Stone T, et al. Uncertainty quantification of reference-based cellular deconvolution algorithms. Epigenetics. 2023;18(1):2137659.

56. Gorrie-Stone TJ, Smart MC, Saffari A, Malki K, Hannon E, Burrage J, et al. Bigmelon: tools for analysing large DNA methylation datasets. Bioinformatics. 2019;35(6):981–6.

57. Pidsley R, Y Wong CC, Volta M, Lunnon K, Mill J, Schalkwyk LC. A data-driven approach to preprocessing Illumina 450K methylation array data. BMC Genomics. 2013;14:293.

58. Chen YA, Lemire M, Choufani S, Butcher DT, Grafodatskaya D, Zanke BW, et al. Discovery of cross-reactive probes and polymorphic CpGs in the Illumina Infinium HumanMethylation450 microarray. Epigenetics. 2013;8(2):203–9.

59. Price ME, Cotton AM, Lam LL, Farré P, Emberly E, Brown CJ, et al. Additional annotation enhances potential for biologically-relevant analysis of the Illumina Infinium HumanMethylation450 BeadChip array. Epigenetics Chromatin. 2013;6(1):4.

60. Aryee MJ, Jaffe AE, Corrada-Bravo H, Ladd-Acosta C, Feinberg AP, Hansen KD, et al. Minfi: a flexible and comprehensive Bioconductor package for the analysis of Infinium DNA methylation microarrays. Bioinformatics. 2014;30(10):1363–9.

61. Pidsley R, Wong CCY, Volta M, Lunnon K, Mill J, Schalkwyk LC. A data-driven approach to preprocessing Illumina 450K methylation array data. Bmc Genomics. 2013;14.

62. Schwarzer G. meta: An R Package for meta-analysis. R News. 2007;7:40–5.

