## Supplementary Figure for "Quantifying the proportion of different cell types in the human cortex using DNA methylation profiles"

**Supplementary Figure 1**. Schematic of gating strategies for isolate brain cell types. Panels A,B,C are strategies used by the Complex Disease Epigenetic Group at Exeter, with panel D represents the strategy used to isolate cell types in the EPIGABA project. The blue boxes state the combination of antibodies used and the white boxes give the labels used to refer to that sample type in the manuscript.

**B**

**D**

**C**

**A**

Prefrontal Cortex

NeuN+ neuronal enriched

NeuN- glial enriched

NeuN-/SOX10+ oligodendrocyte enriched

NeuN-/SOX10- mircoglial & astrocyte enriched

NeuN-/SOX10-/IRF8+ microglial enriched

NeuN-/SOX10-/IRF8- astrocyte enriched

Prefrontal Cortex

NeuN+ neuronal enriched

NeuN- glial enriched

NeuN-/SOX10+ oligodendrocyte enriched

NeuN-/SOX10- mircoglial & astrocyte enriched

Occipital Cortex

NeuN+ neuronal enriched

NeuN+/SOX6+ gabaergic neuronal enriched

NeuN+/SOX6- gluamatergic neuronal enriched

NeuN- glial enriched

Prefrontal Cortex

SATB2+ inhibatory neuronal enriched

SATB2- exhibatory neuronal & glial enriched

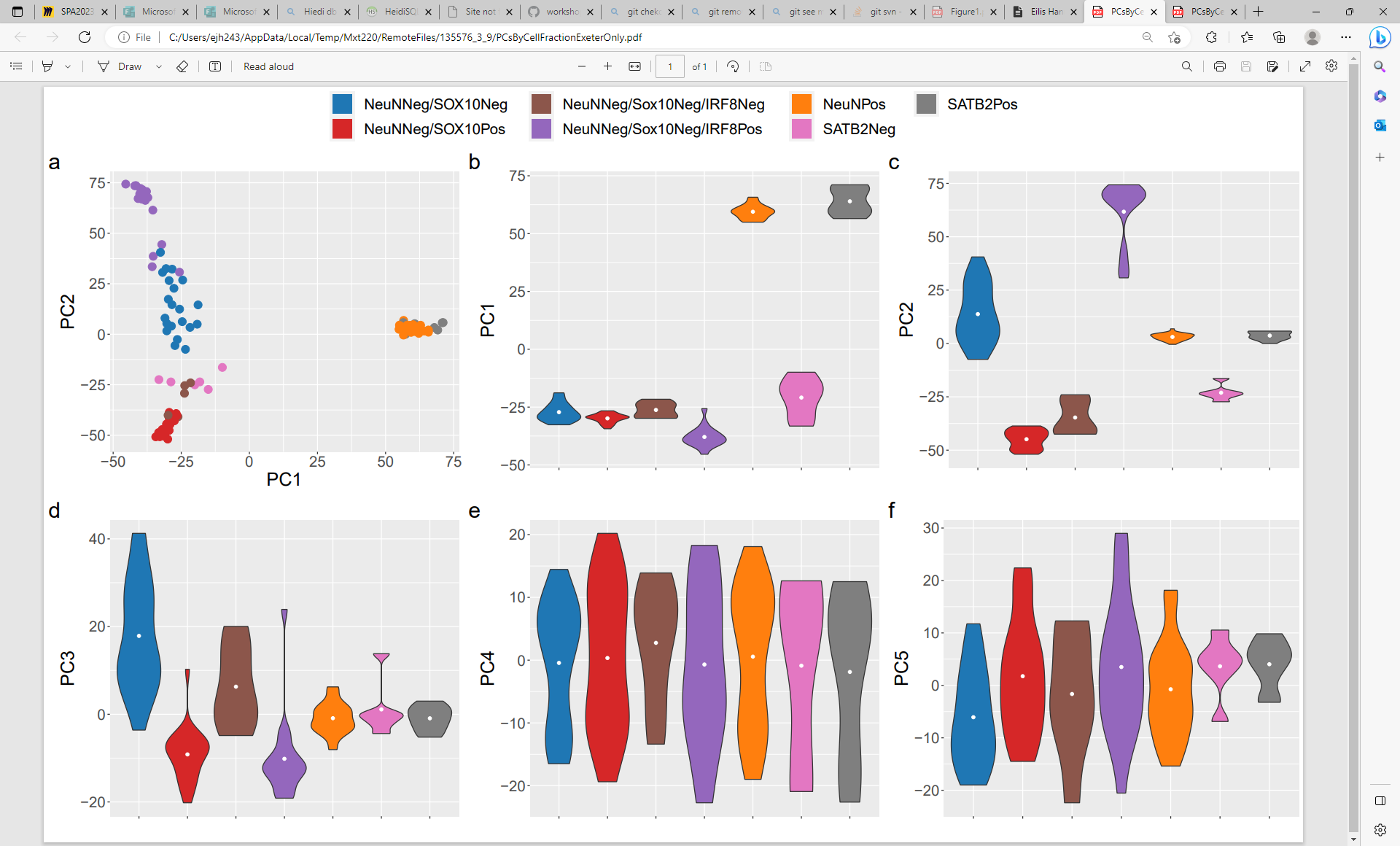

**Supplementary Figure 2.** **Major axes of variation in DNA methylation data are driven by cell type**. Scatterplot (a) of the first two principal components (first, second and sixth) that optimally separate different sample types. Each point represents a sample and the colour of the point indicates the nuclei fraction. Violin plots (b-f) for the first 5 principal components grouped by nuclei fraction.

**Supplementary Figure 3.** Schematic of reference panels form computational deconvolution of cellular composition in brain tissue. Each row represents a separate reference panel with a different combination of cell types for which deconvolution was performed.

NeuNPos

NeuNNeg/Sox10Pos

NeuNNeg/Sox10Neg

NeuNPos

NeuNNeg/Sox10Pos

NeuNNeg/Sox10Neg/IRF8Pos

NeuNNeg/Sox10Neg

/IRF8Neg

SATB2Pos

SATB2Neg

NeuNPos

NeuNNeg/Sox10Pos

NeuNNeg/Sox10Neg

NeuNPos

NeuNNeg/Sox10Pos

NeuNNeg/Sox10Neg/IRF8Pos

NeuNNeg/Sox10Neg

/IRF8Neg

SATB2Pos

SATB2Pos

NeuNPos

/SOX6Pos

NeuNPos

/SOX6Neg

NeuNNeg

NeuNPos

/SOX6Pos

NeuNPos

/SOX6Neg

NeuNPos

/SOX6Pos

NeuNPos

/SOX6Neg

NeuNNeg/Sox10Pos

NeuNNeg/Sox10Neg

NeuNNeg/Sox10Pos

NeuNNeg/Sox10Neg

/IRF8Pos

1

2

3

4

5

6

7

8

NeuNNeg/Sox10Neg

/IRF8Neg

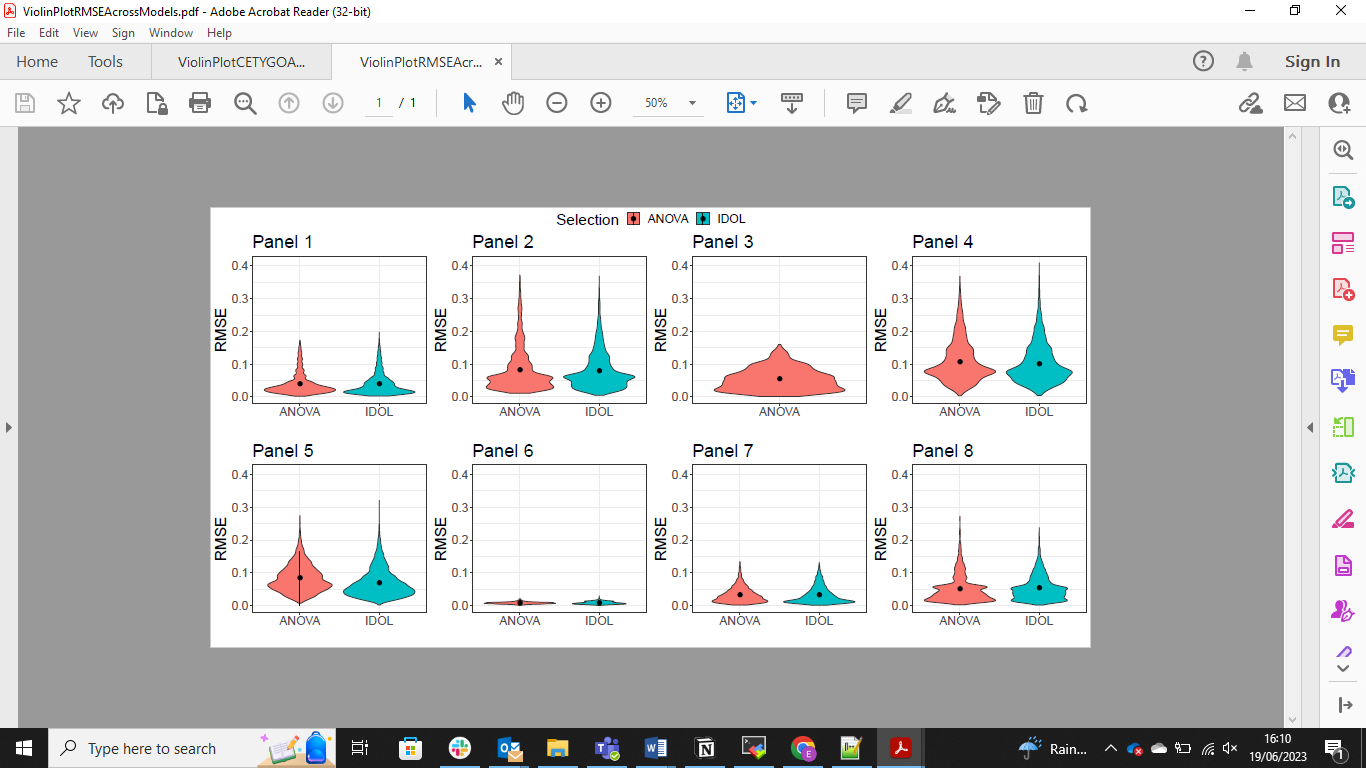

**Supplementary Figure 4.** Violin plots of the root mean square error (RMSE) between the fixed cellular proportions used to construct the bulk brain profiles and the estimated proportions. Each figure panel represents a different panel of cell types (**Table 1**/**Supplementary Figure 3**).

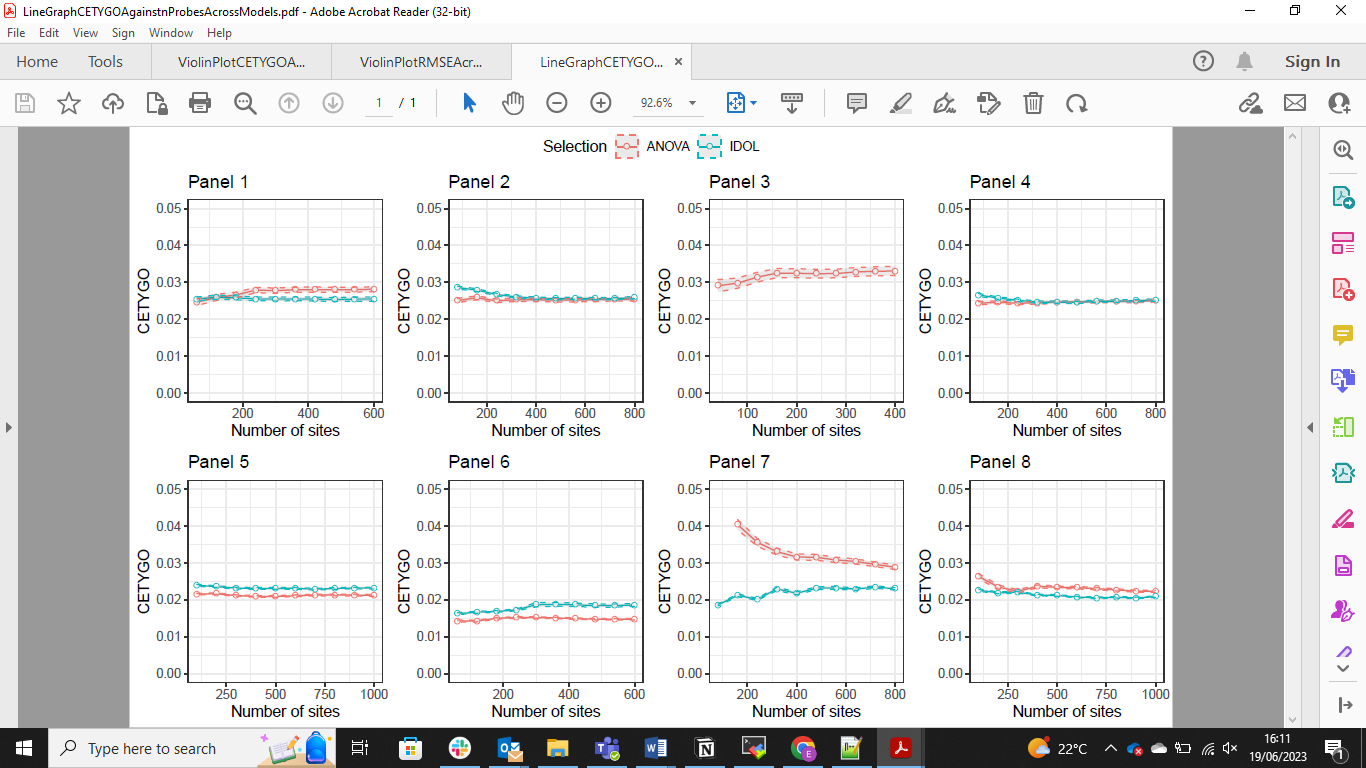

**Supplementary Figure 5.** Line graph of the effect of number of cell-specific probes on the accuracy of cellular deconvolution measured by the CETYGO score. Each figure panel represents a different panel of cell types (**Table 1**/**Supplementary Figure 3**).

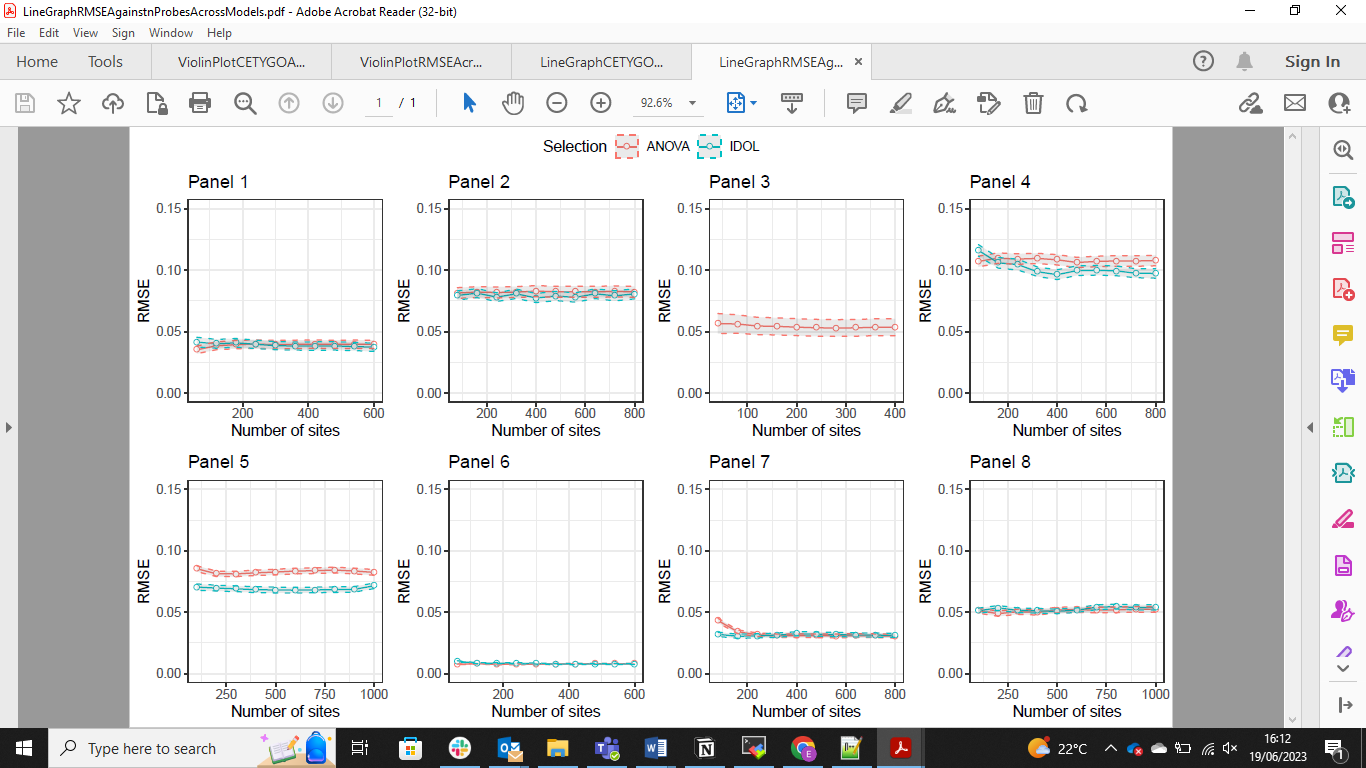

**Supplementary Figure 6.** Line graph of the effect of number of cell-specific probes on the accuracy of cellular deconvolution measured by the root mean square error (RMSE) between the fixed cellular proportions used to construct the bulk brain profiles and the estimated proportions. Each figure panel represents a different panel of cell types (**Table 1**/**Supplementary Figure 3**).

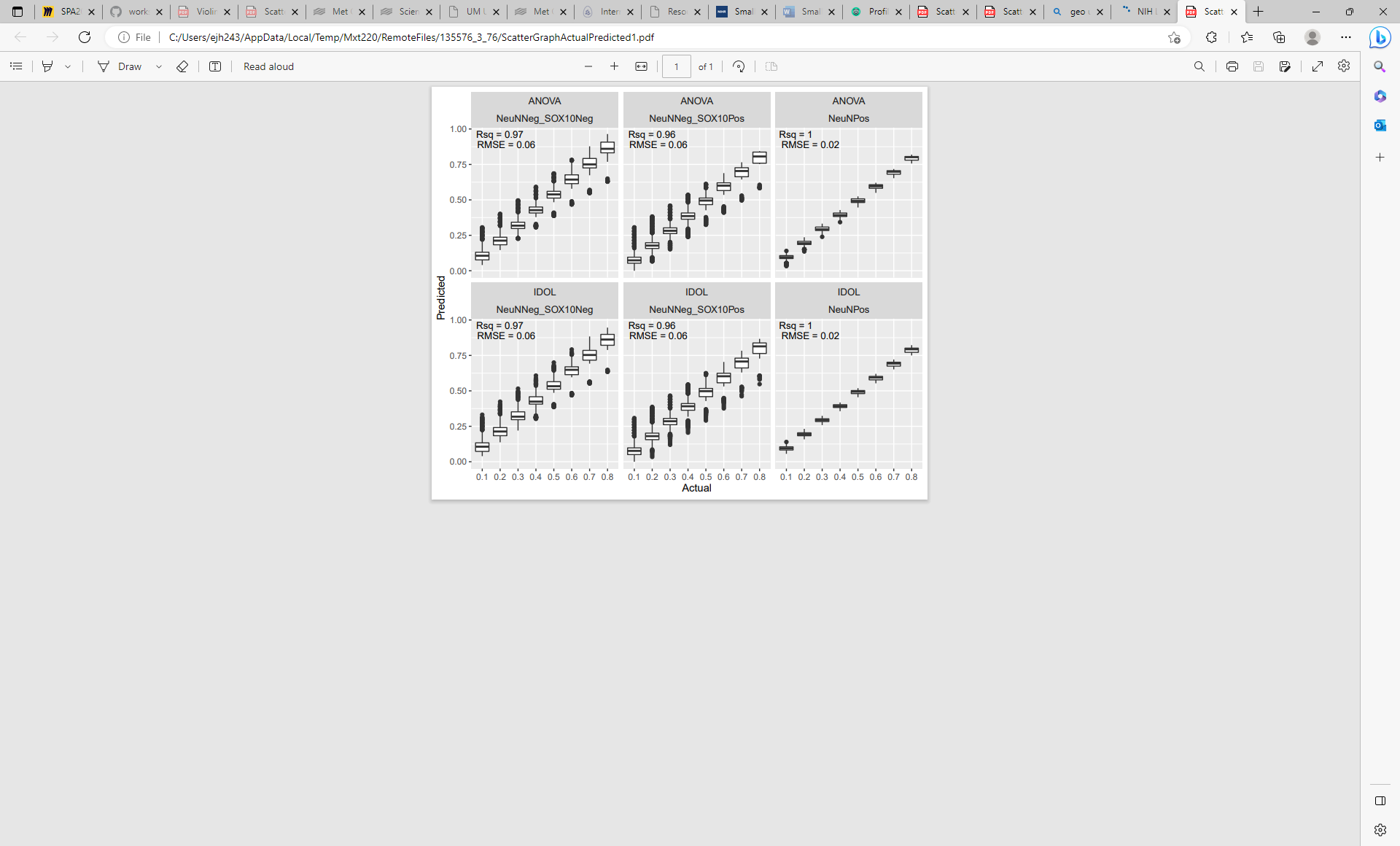

**Supplementary Figure 7.** Boxplots of distribution of predicted proportion (y-axis), grouped by actual proportion specified in the reconstruction of brain profiles (x-axis). Shown here are the results for panel 1 of cell types (**Table 1**/**Supplementary Figure 3**) using both the ANOVA and IDOL methods to select cell specific sites. Each panel in the figure represents a different cell type.

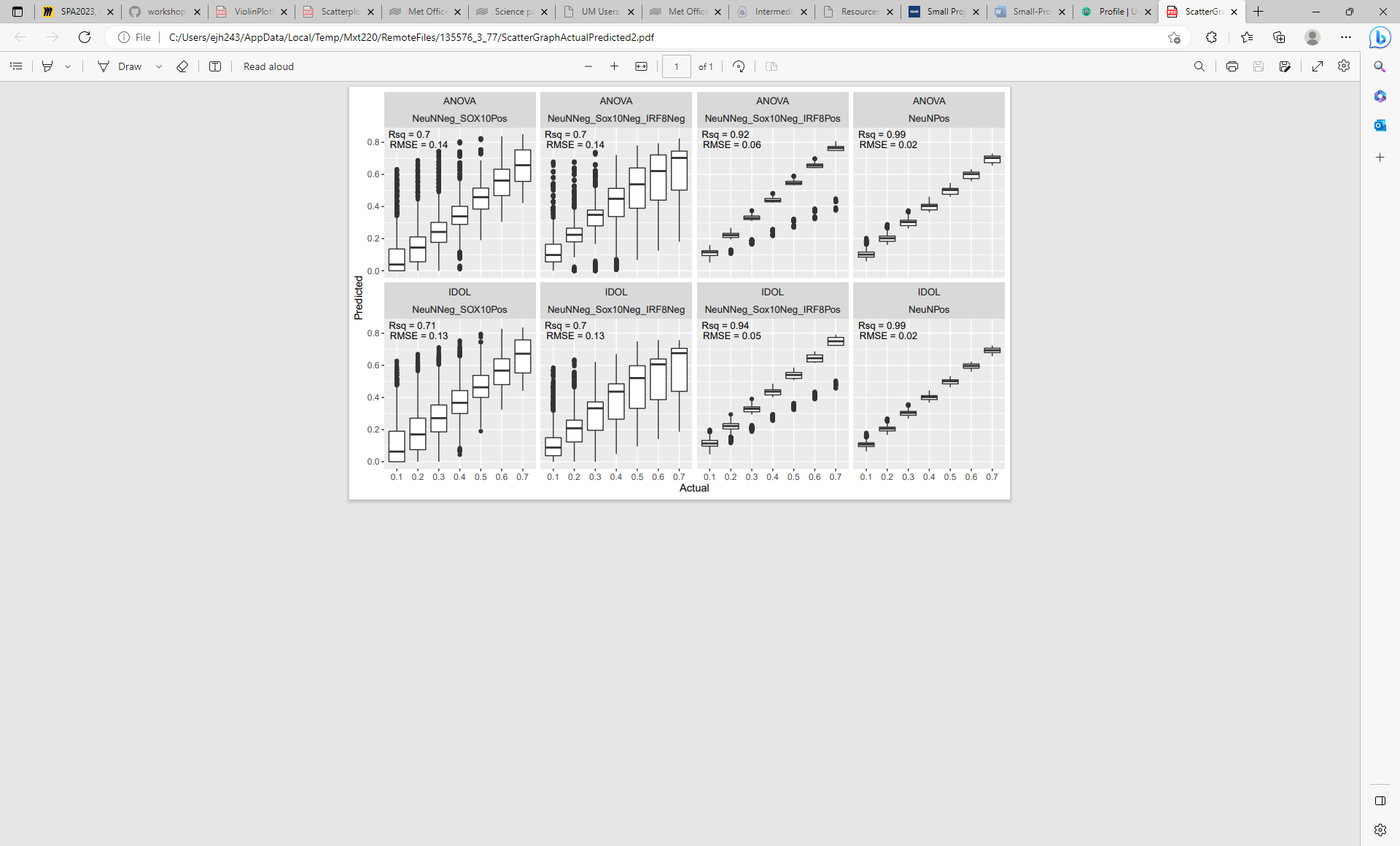

**Supplementary Figure 8** .Boxplots of distribution of predicted proportion (y-axis), grouped by actual proportion specified in the reconstruction of brain profiles (x-axis). Shown here are the results for panel 2 of cell types (**Table 1**/**Supplementary Figure 3**) using both the ANOVA and IDOL methods to select cell specific sites. Each panel in the figure represents a different cell type.

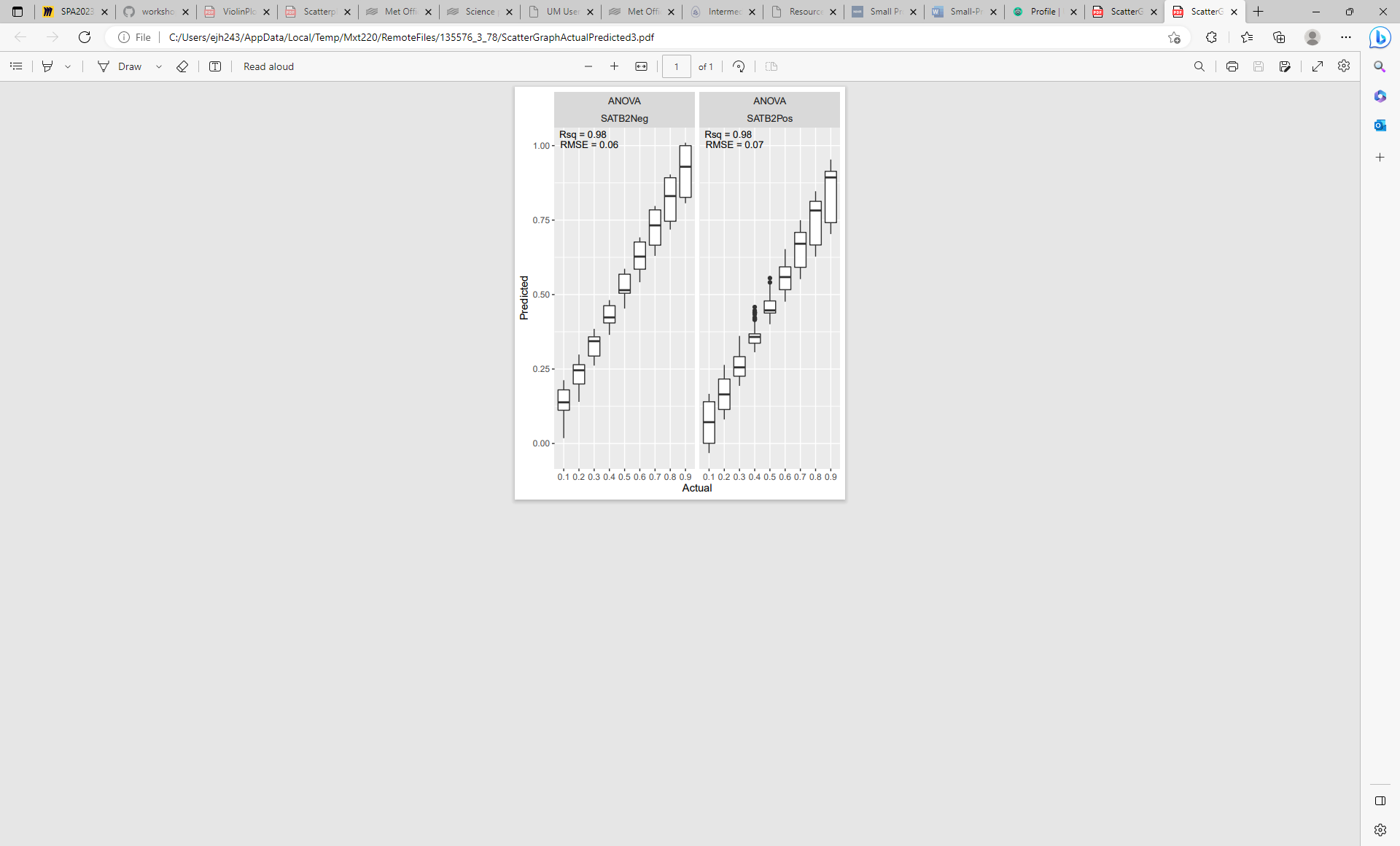

**Supplementary Figure 9.** Boxplots of distribution of predicted proportion (y-axis), grouped by actual proportion specified in the reconstruction of brain profiles (x-axis). Shown here are the results for panel 3 of cell types (**Table 1**/**Supplementary Figure 3**) using both the ANOVA method to select cell specific sites. Each panel in the figure represents a different cell type.

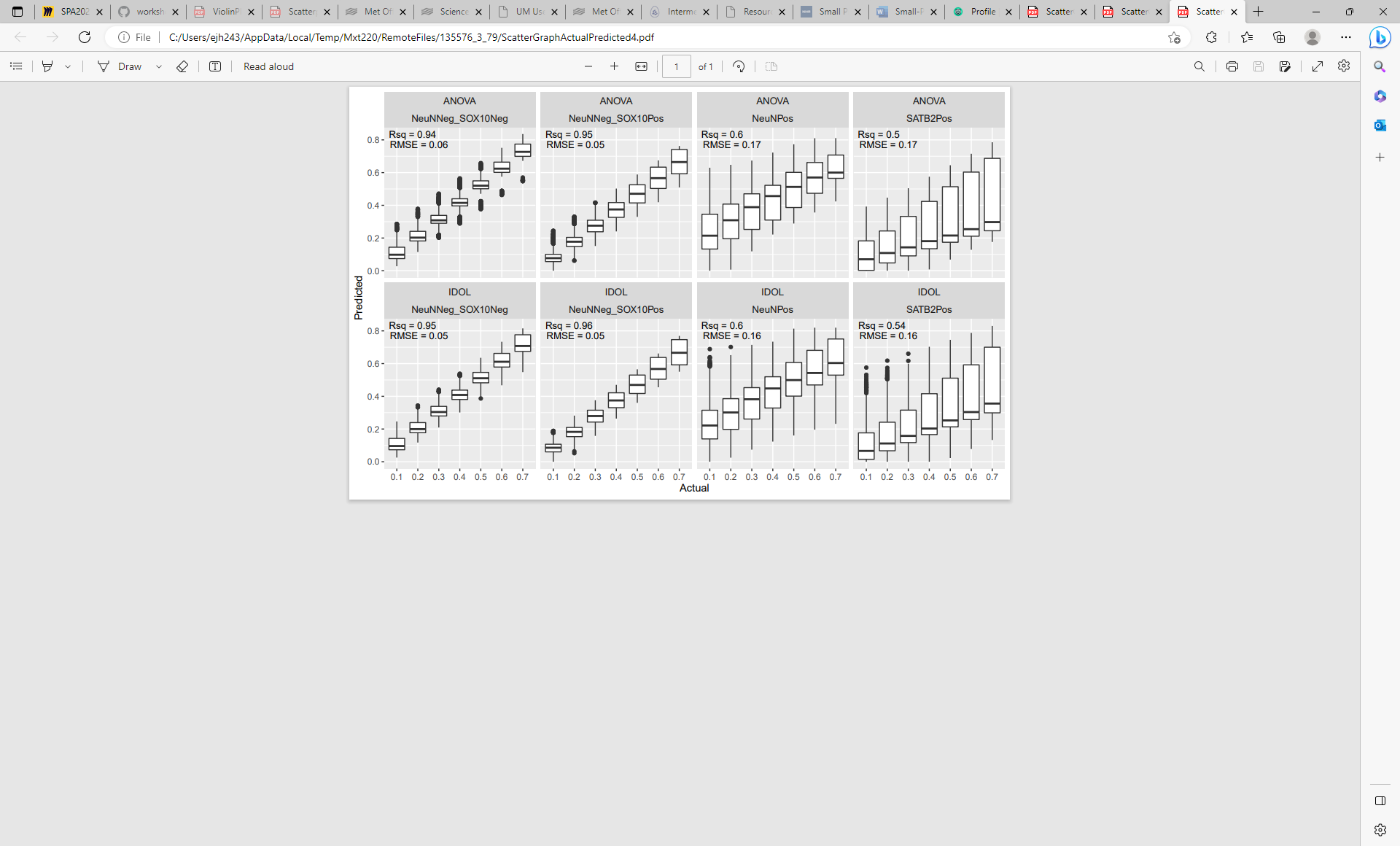

**Supplementary Figure 10.** Boxplots of distribution of predicted proportion (y-axis), grouped by actual proportion specified in the reconstruction of brain profiles (x-axis). Shown here are the results for panel 4 of cell types (**Table 1**/**Supplementary Figure 3**) using both the ANOVA and IDOL methods to select cell specific sites. Each panel in the figure represents a different cell type.

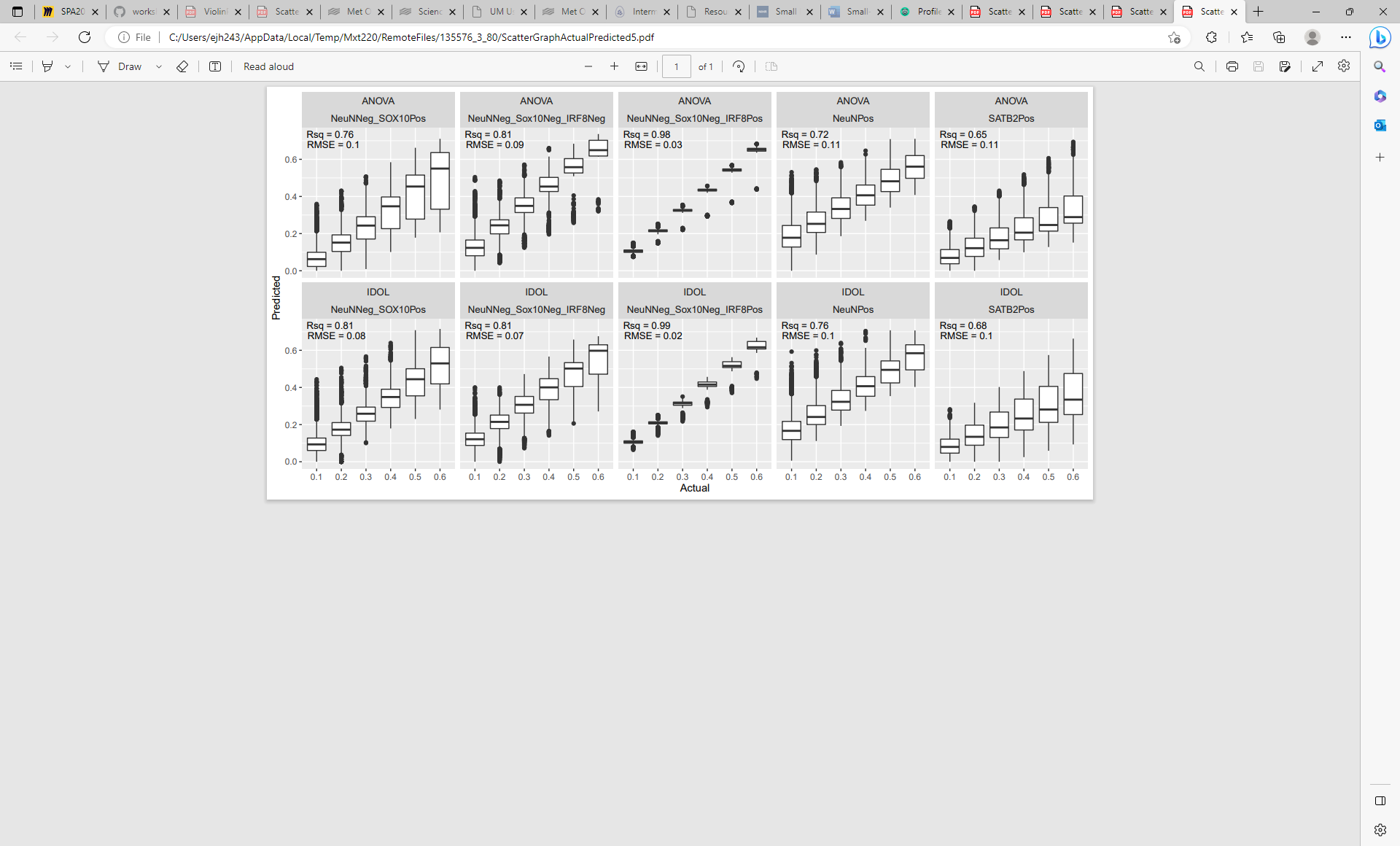

**Supplementary Figure 11.** Boxplots of distribution of predicted proportion (y-axis), grouped by actual proportion specified in the reconstruction of brain profiles (x-axis). Shown here are the results for panel 5 of cell types (**Table 1**/**Supplementary Figure 3**) using both the ANOVA and IDOL methods to select cell specific sites. Each panel in the figure represents a different cell type.

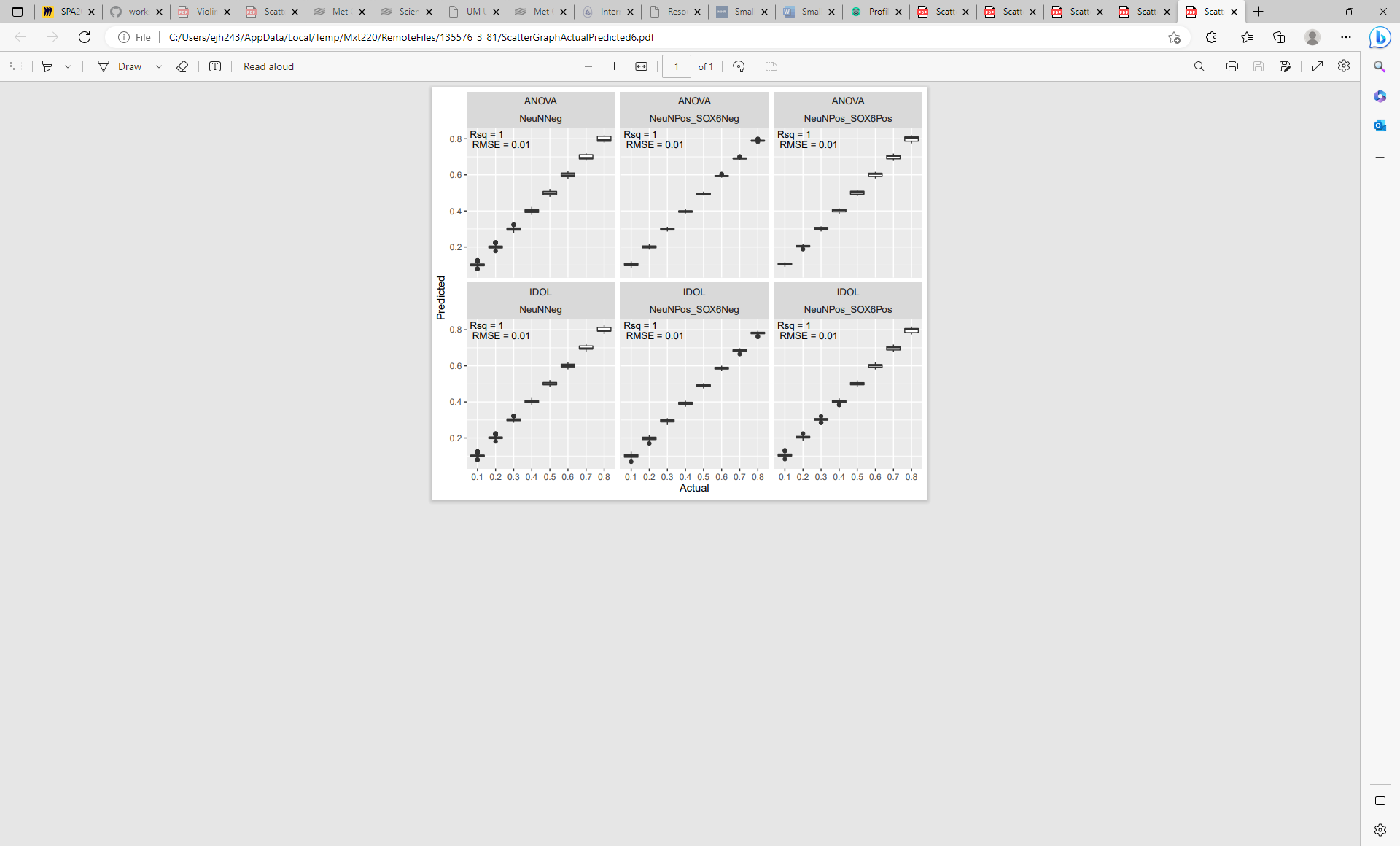

**Supplementary Figure 12.** Boxplots of distribution of predicted proportion (y-axis), grouped by actual proportion specified in the reconstruction of brain profiles (x-axis). Shown here are the results for panel 6 of cell types (**Table 1**/**Supplementary Figure 3**) using both the ANOVA and IDOL methods to select cell specific sites. Each panel in the figure represents a different cell type.

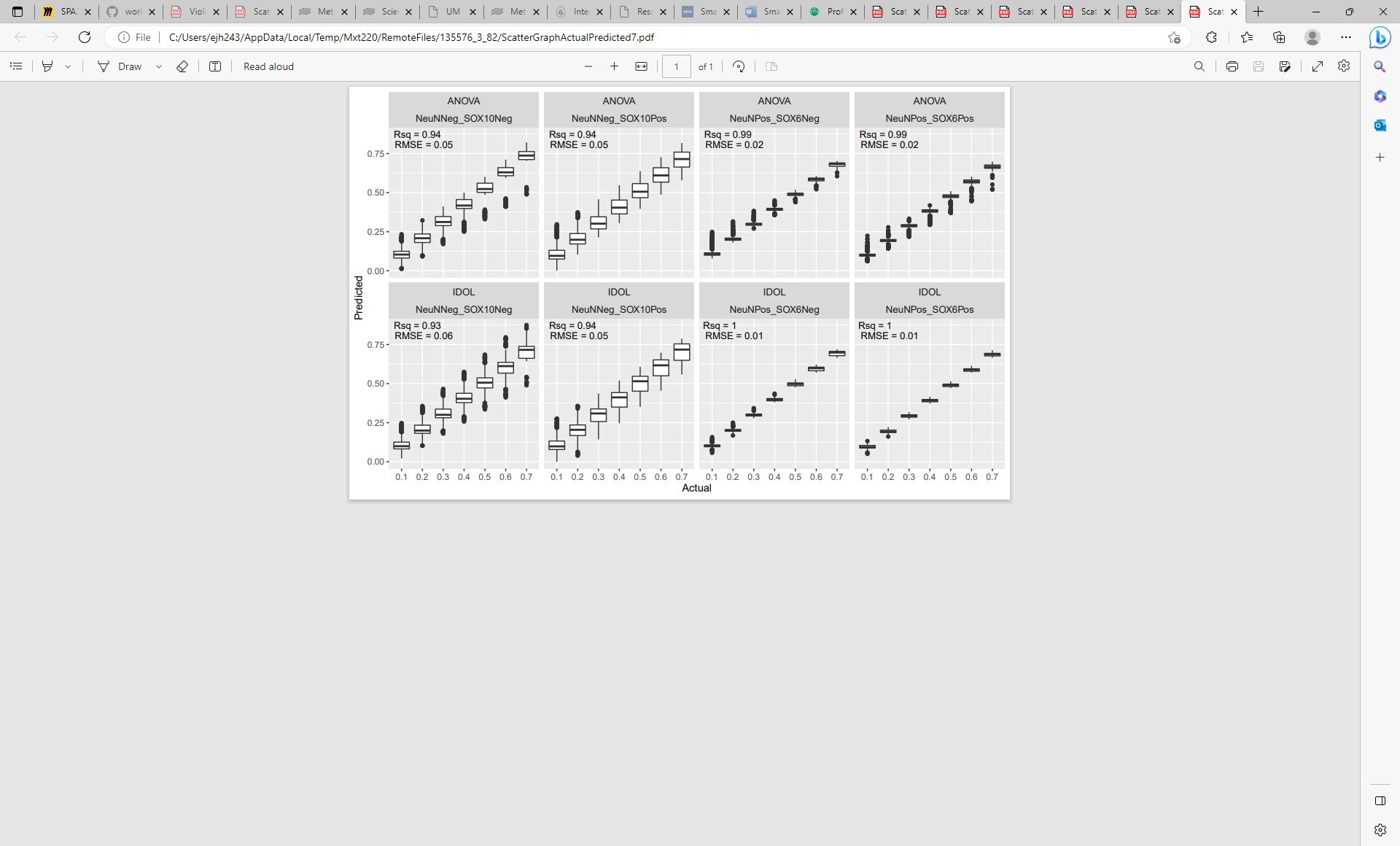

**Supplementary Figure 13.** Boxplots of distribution of predicted proportion (y-axis), grouped by actual proportion specified in the reconstruction of brain profiles (x-axis). Shown here are the results for panel 7 of cell types (**Table 1**/**Supplementary Figure 3**) using both the ANOVA and IDOL methods to select cell specific sites. Each panel in the figure represents a different cell type.

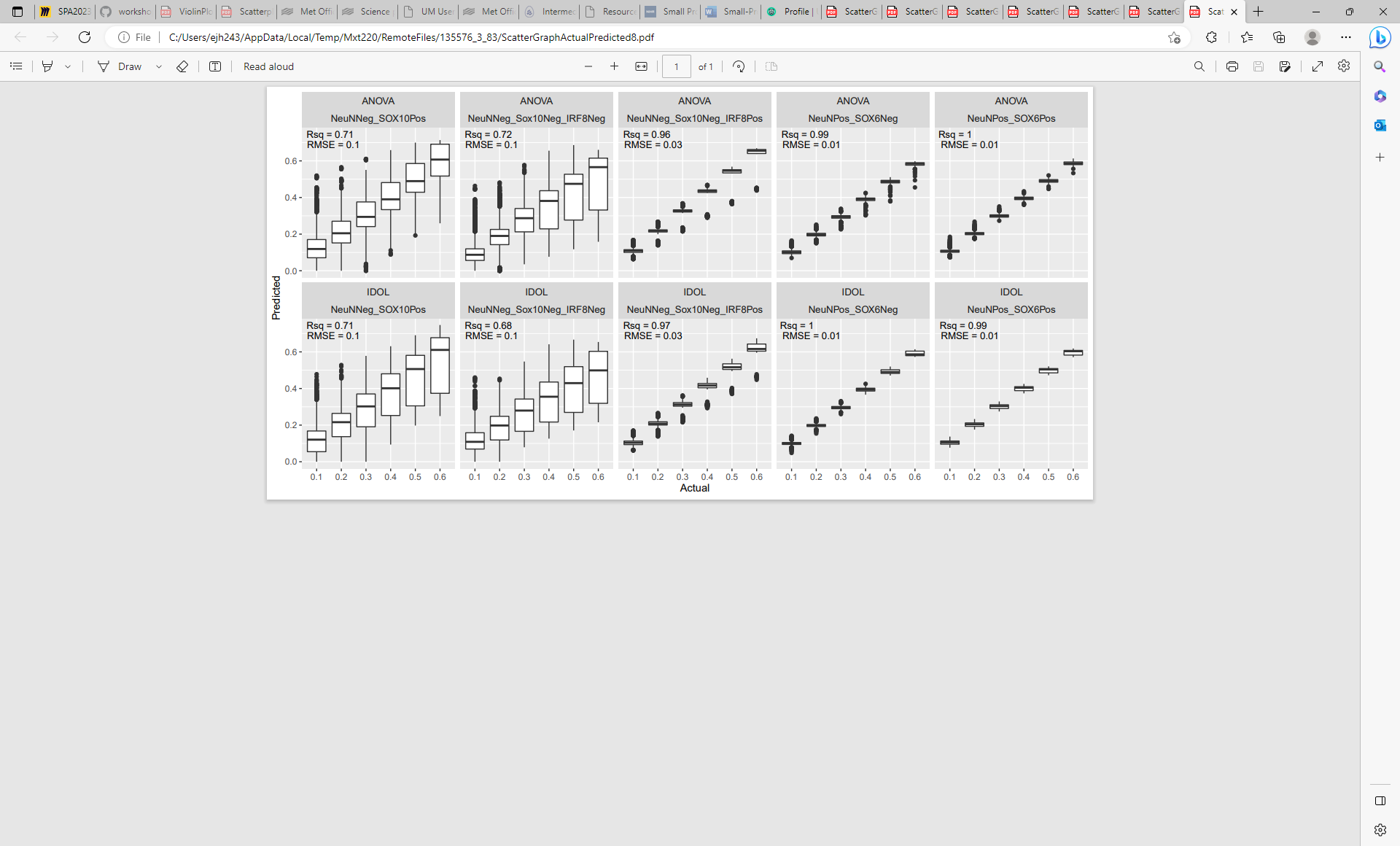

**Supplementary Figure 14.** Boxplots of distribution of predicted proportion (y-axis), grouped by actual proportion specified in the reconstruction of brain profiles (x-axis). Shown here are the results for panel 8 of cell types (**Table 1**/**Supplementary Figure 3**) using both the ANOVA and IDOL methods to select cell specific sites. Each panel in the figure represents a different cell type.

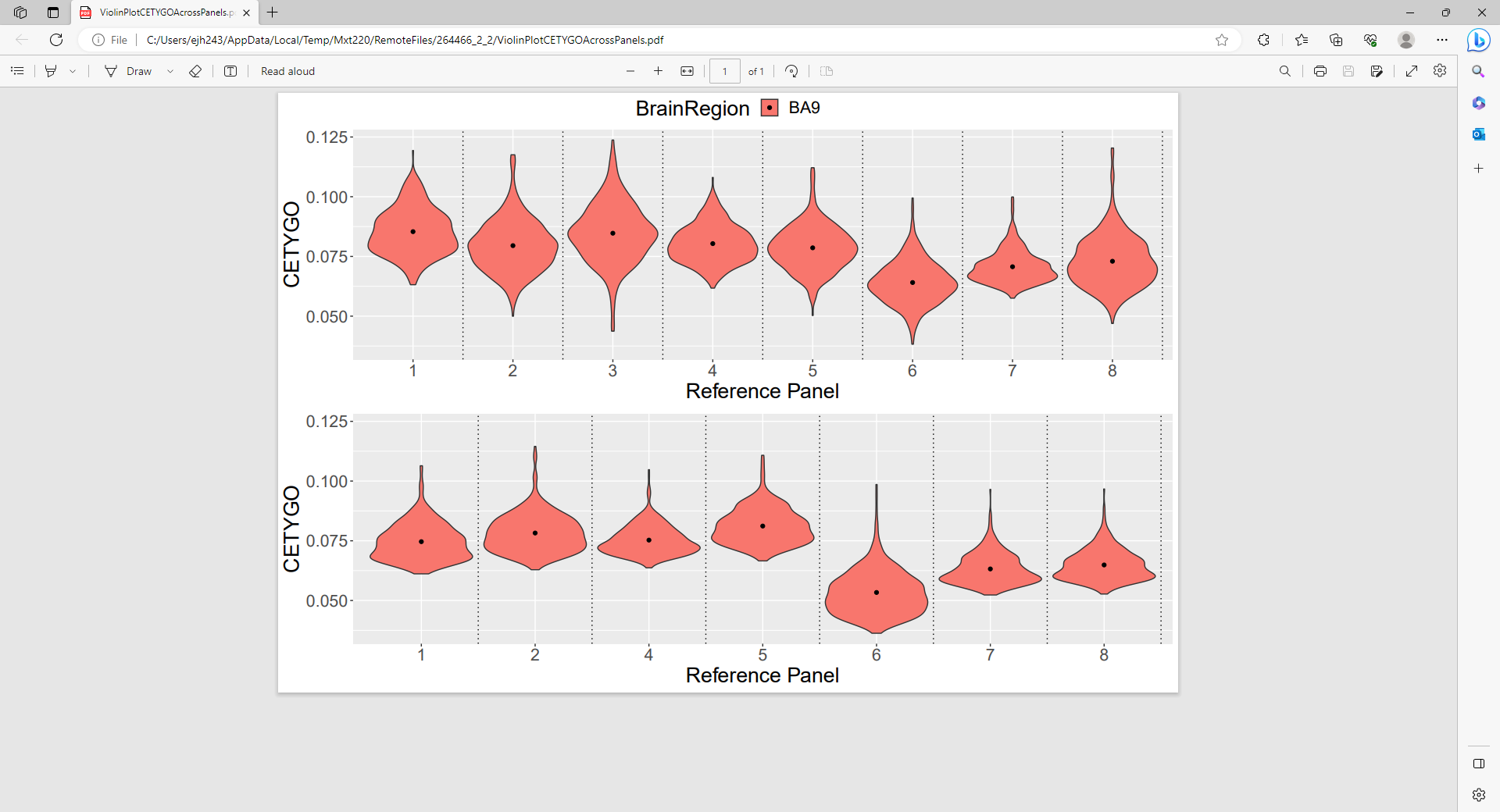

**Supplementary Figure 15A**. Violin plot of CETYGO score across 377 adult prefrontal cortex samples. Top panel, probes cell specific sites were selected with an ANOVA, bottom panel, cell specific sites were selected with the IDOL method.

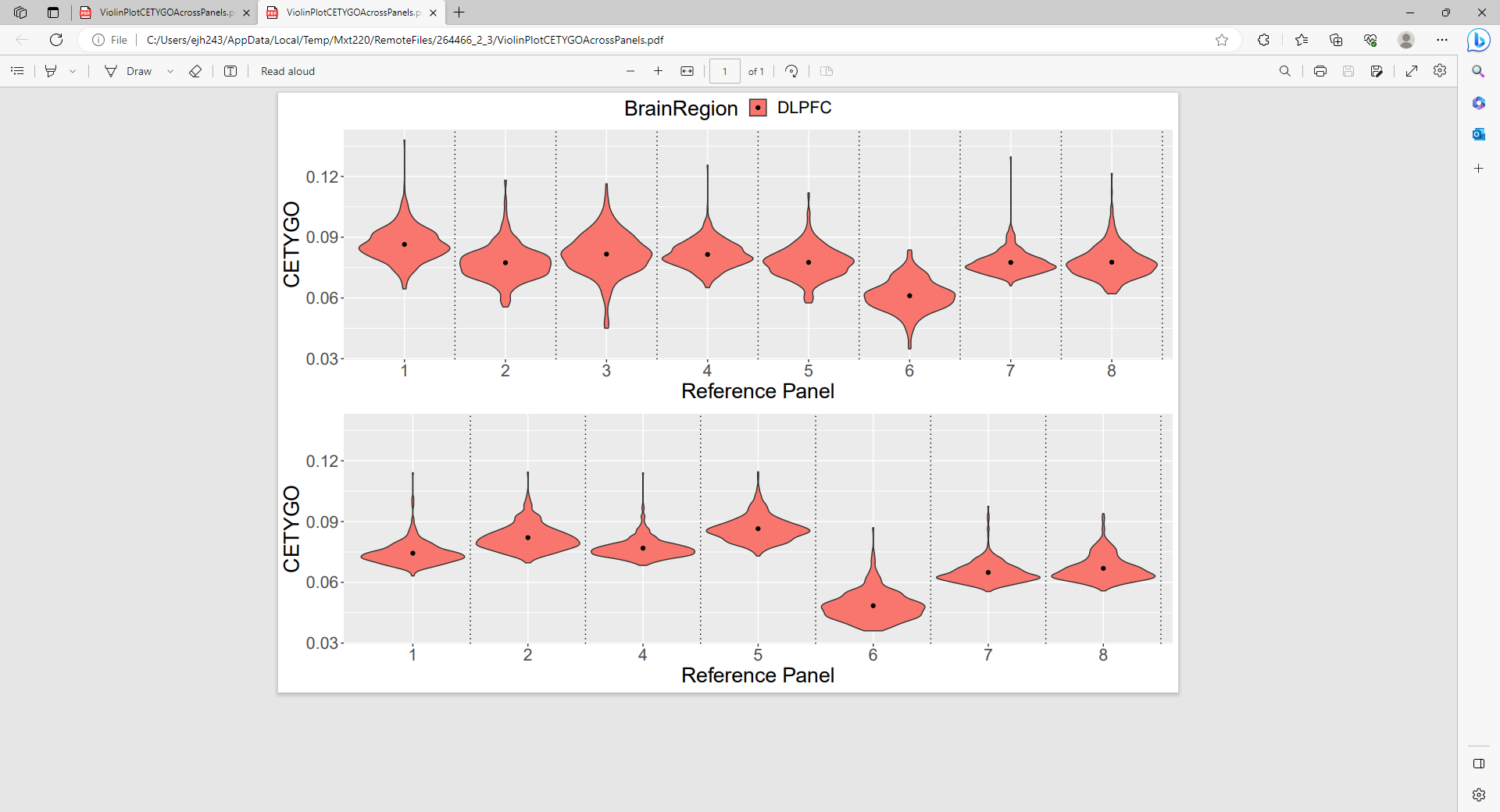

**Supplementary Figure 15B**. Violin plot of CETYGO score across 415 adult prefrontal cortex samples from the Jaffe dataset. Top panel, probes cell specific sites were selected with an ANOVA, bottom panel, cell specific sites were selected with the IDOL method.

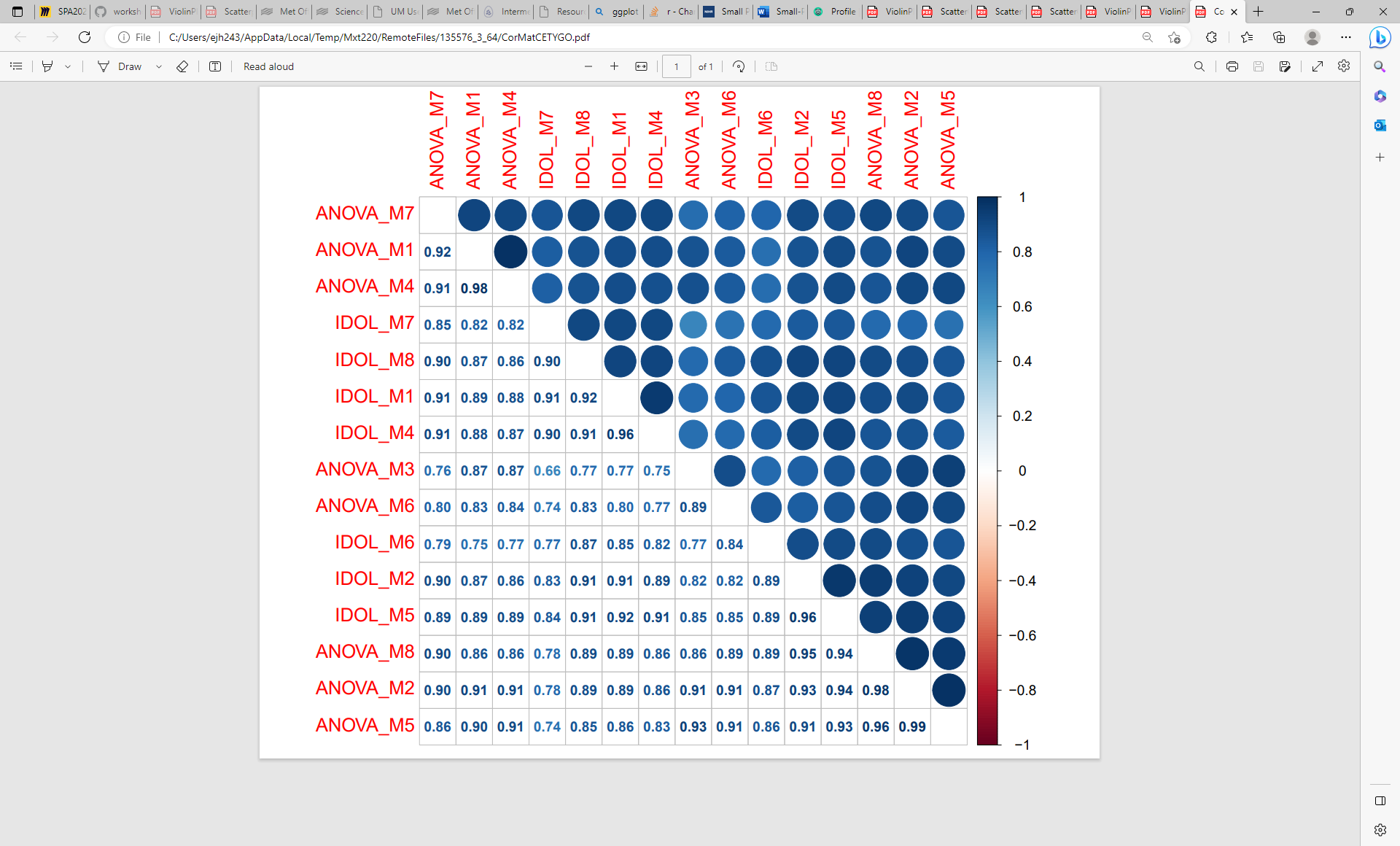

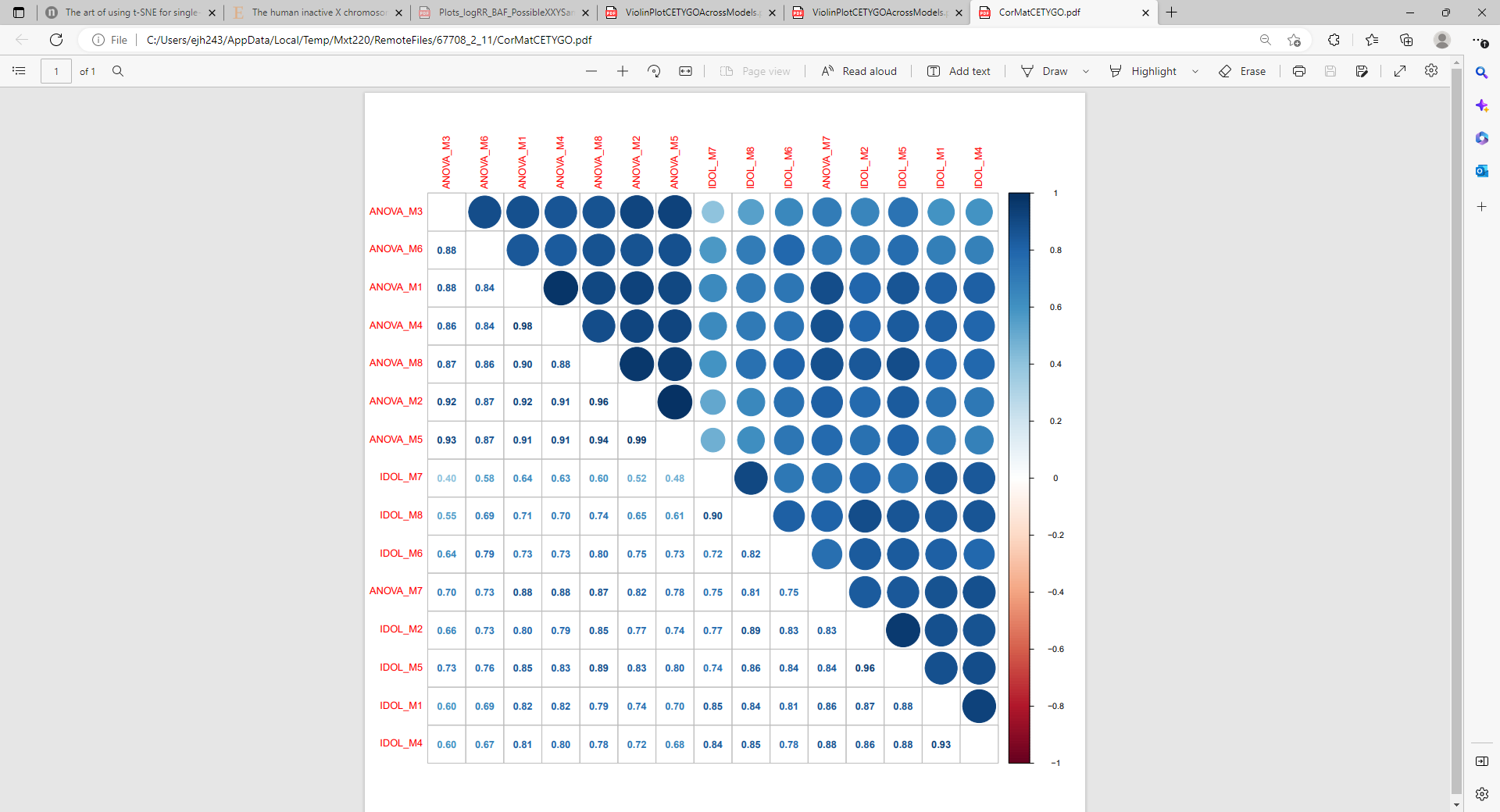

**Supplementary Figure 16.** Correlation heatmap of CETYGO scores from 15 different deconvolution models applied to adult prefrontal samples. Each row/column denotes a different model where the first word (ANOVA or IDOL) indicates how the cell-specific sites were selected and the number after the “M” represent which panel of cell types (as defined in Supplementary Figure 3). The top heatmap is based on the Exeter dataset (n = 377), and the bottom heatmap the Jaffe dataset (n = 415).

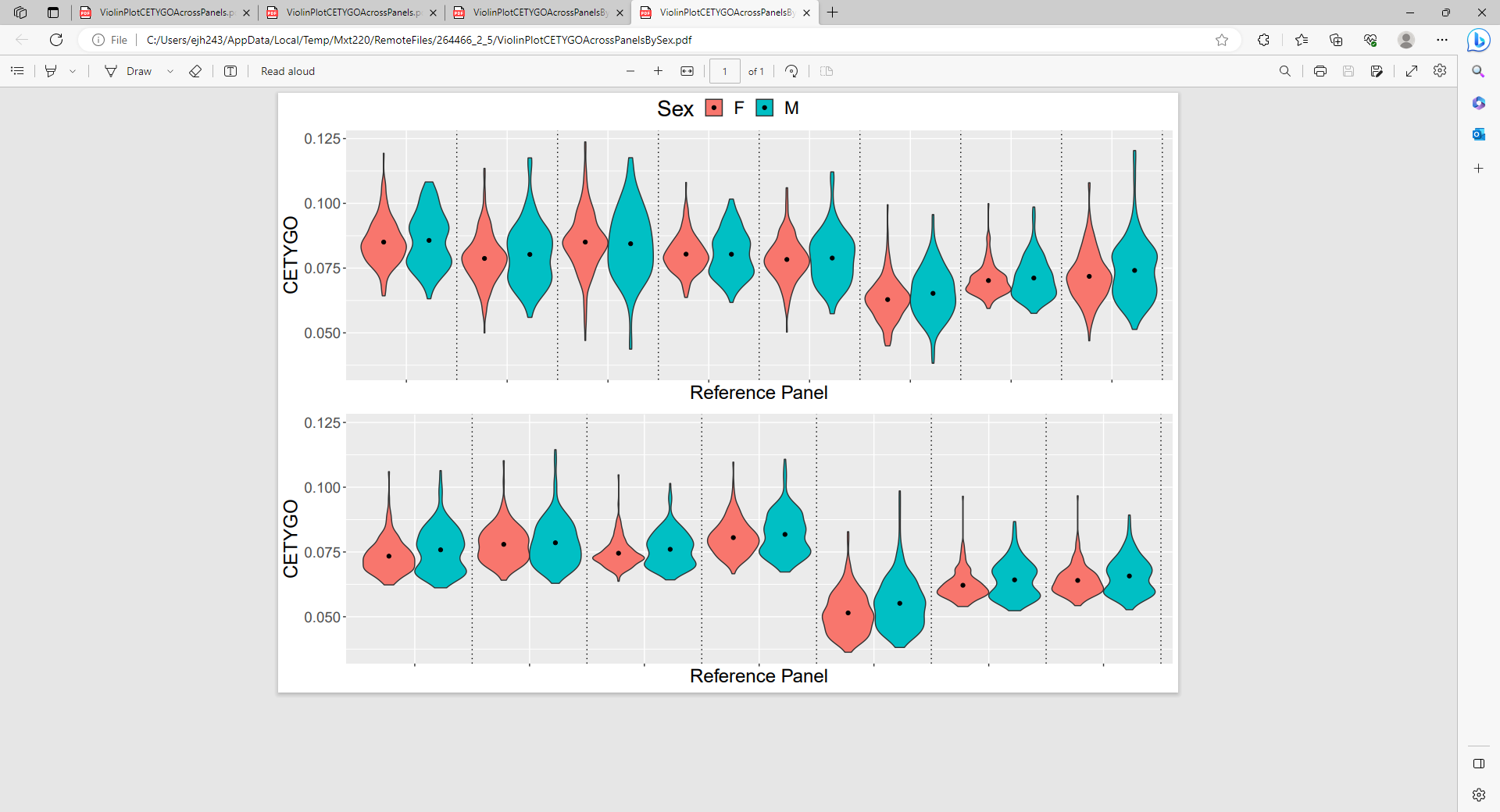

**Supplementary Figure 17A.** Violin plot of CETYGO score grouped by sex across 377 adult prefrontal cortex samples from the Exeter dataset. Top panel, probes cell specific sites were selected with an ANOVA, bottom panel, cell specific sites were selected with the IDOL method.

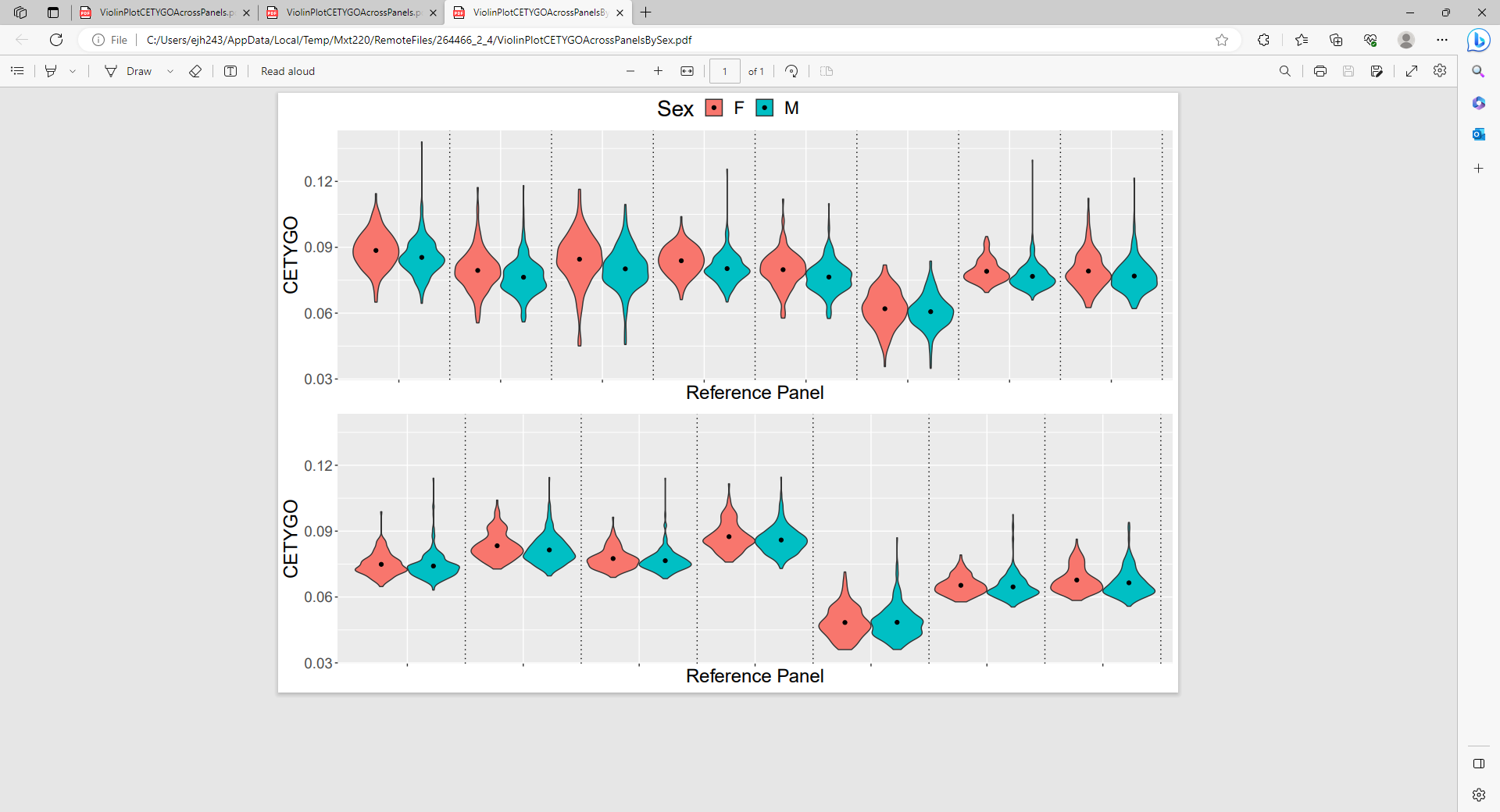

**Supplementary Figure 17B.** Violin plot of CETYGO score grouped by sex across 415 adult prefrontal cortex samples from the Jaffe dataset. Top panel, probes cell specific sites were selected with an ANOVA, bottom panel, cell specific sites were selected with the IDOL method.

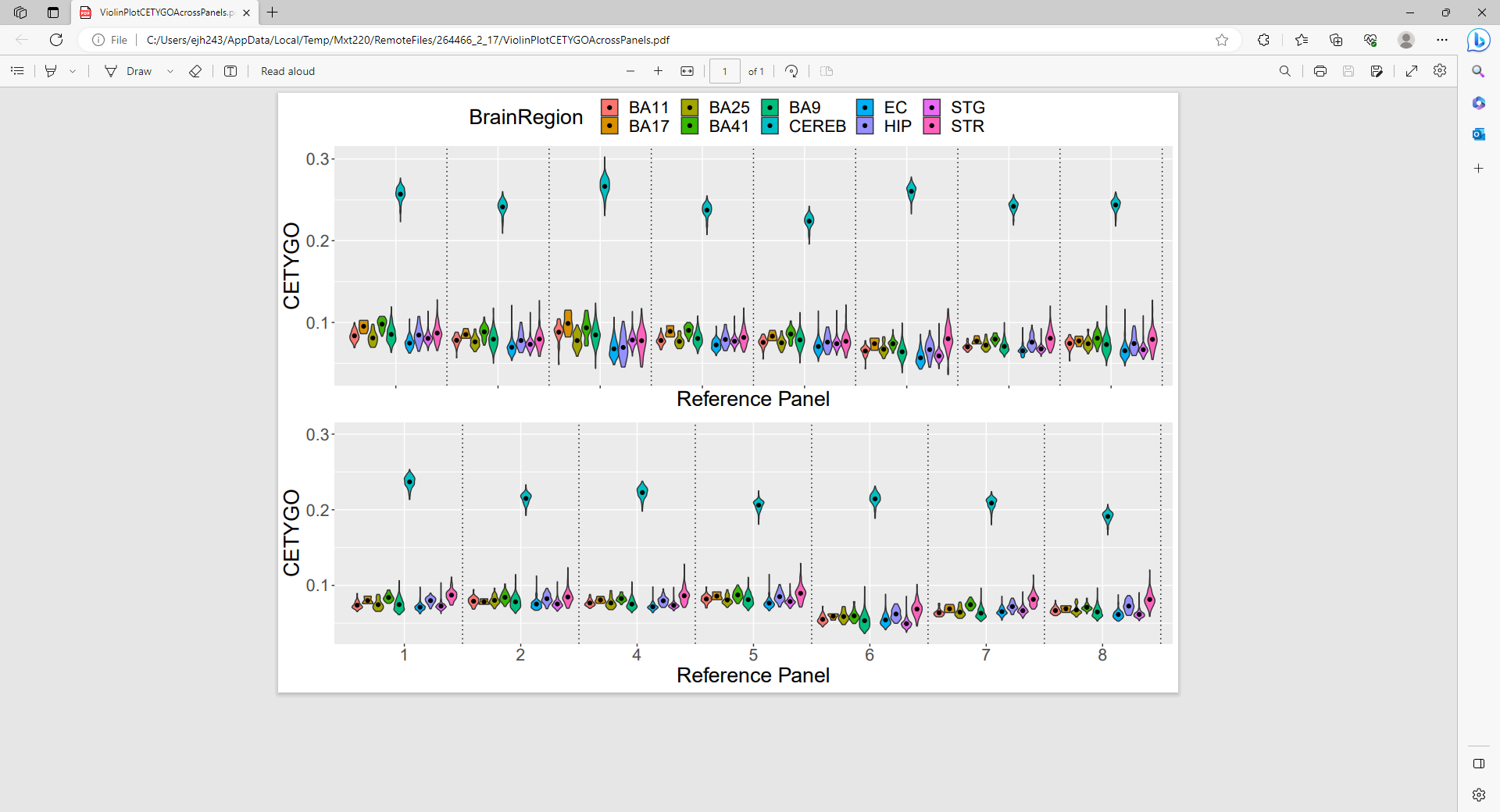

**Supplementary Figure 18.** Violin plot of CETYGO score across 851 adult tissue samples from different brain regions. Top panel, probes cell specific sites were selected with an ANOVA, bottom panel, cell specific sites were selected with the IDOL method.

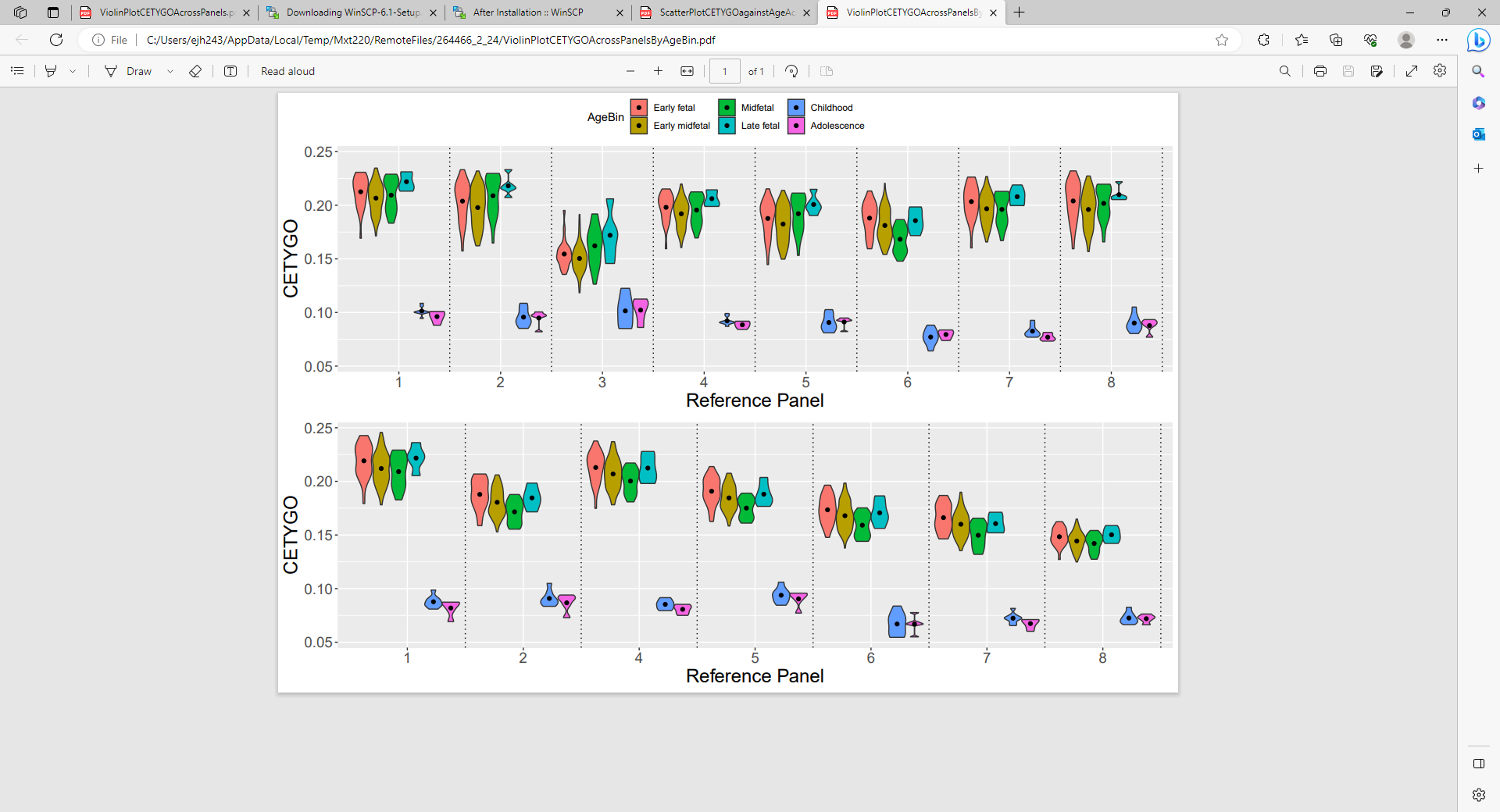

**Supplementary Figure 19.** Violin plot of CETYGO score across 167 fetal and childhood samples grouped into development stages. Top panel, probes cell specific sites were selected with an ANOVA, bottom panel, cell specific sites were selected with the IDOL method.

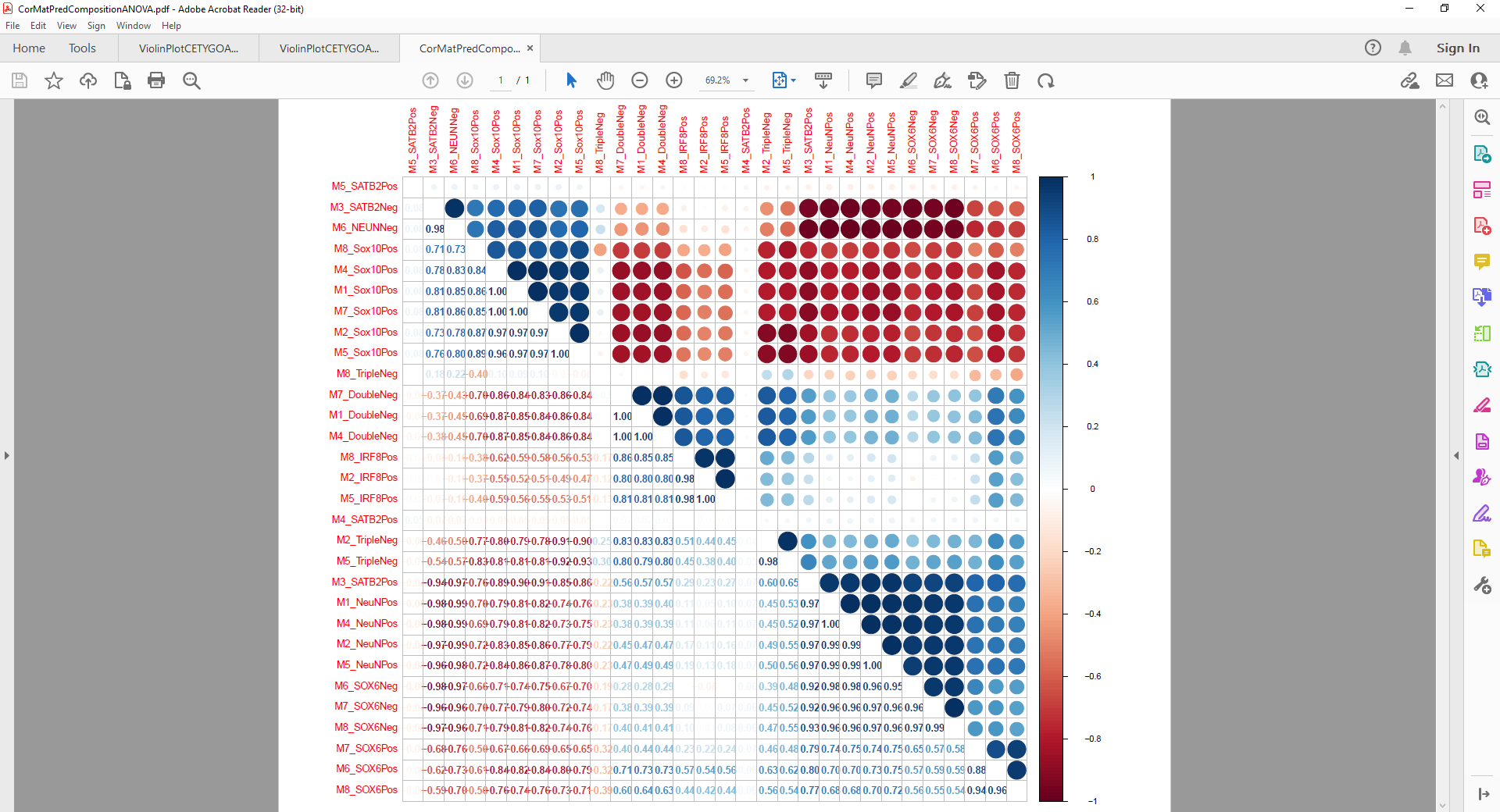

**Supplementary Figure 20.** Heatmap of correlations between predicted cellular proportions across all reference panels with ANOVA to select cell-specific sites in the Exeter dataset (n = 377).

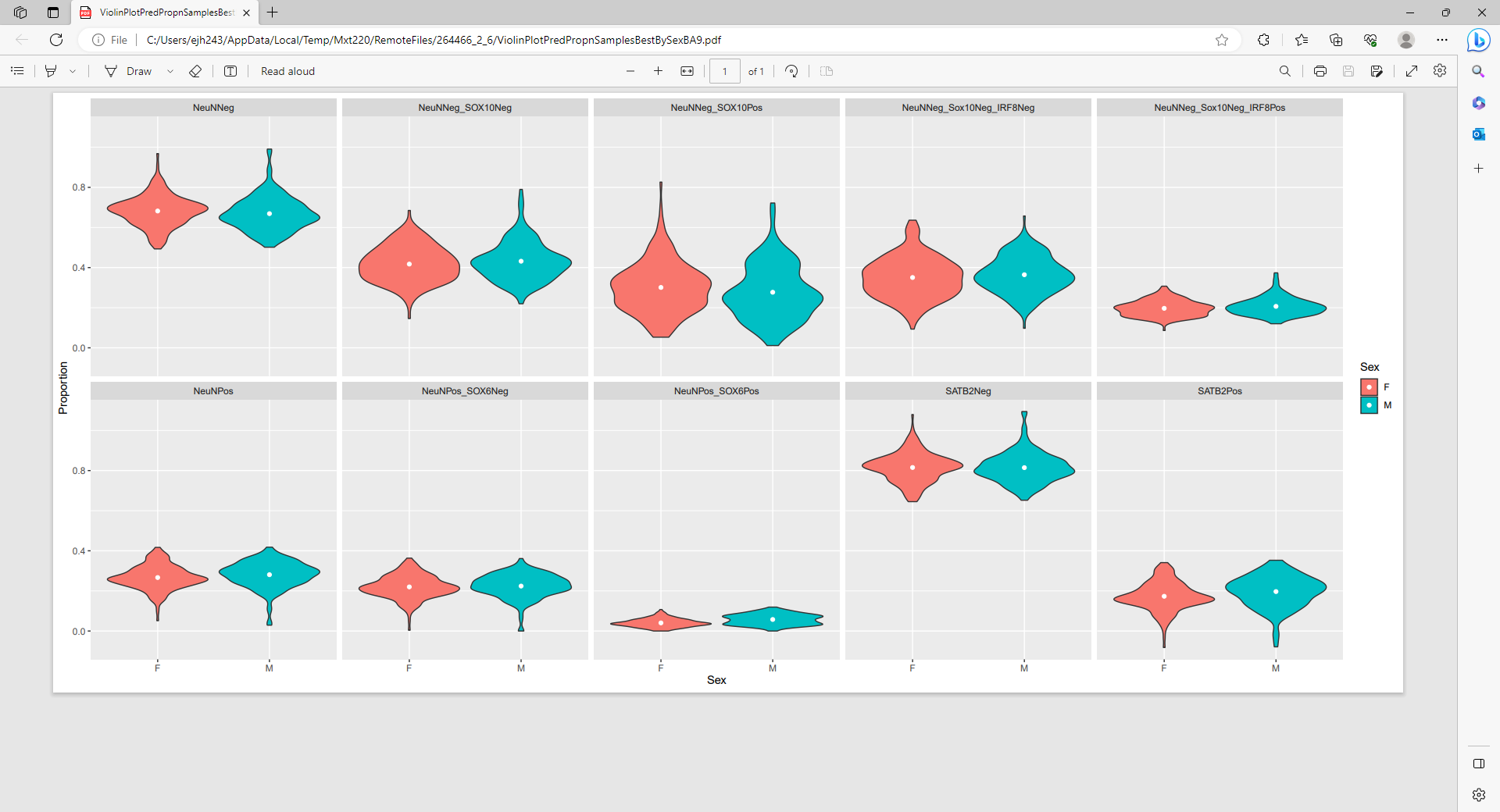

**Supplementary Figure 21.** Violin plots of the distribution of proportion of brain cell types estimated from DNA methylation data generated from the prefrontal cortex grouped by sex across 377 adult prefrontal cortex samples from the Exeter dataset. Each panel represents a different cell type, estimated using the optimal reference panel for that cell type (**Supplementary Table 8**).

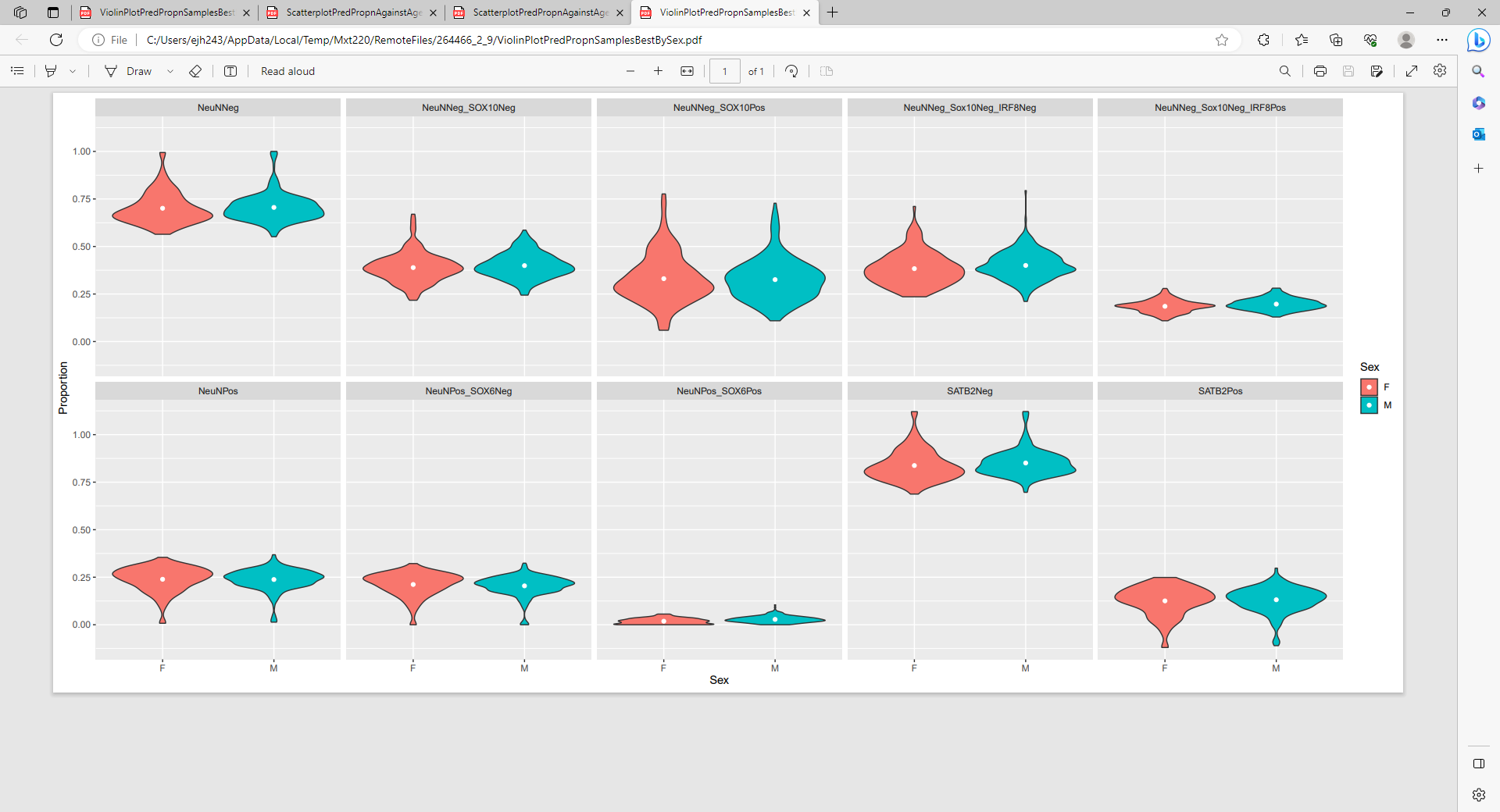

**Supplementary Figure 22.** Violin plots of the distribution of proportion of brain cell types estimated from DNA methylation data generated from the prefrontal cortex grouped by sex across 415 adult prefrontal cortex samples from the Jaffe dataset. Each panel represents a different cell type, estimated using the optimal reference panel for that cell type (**Supplementary Table 8**).

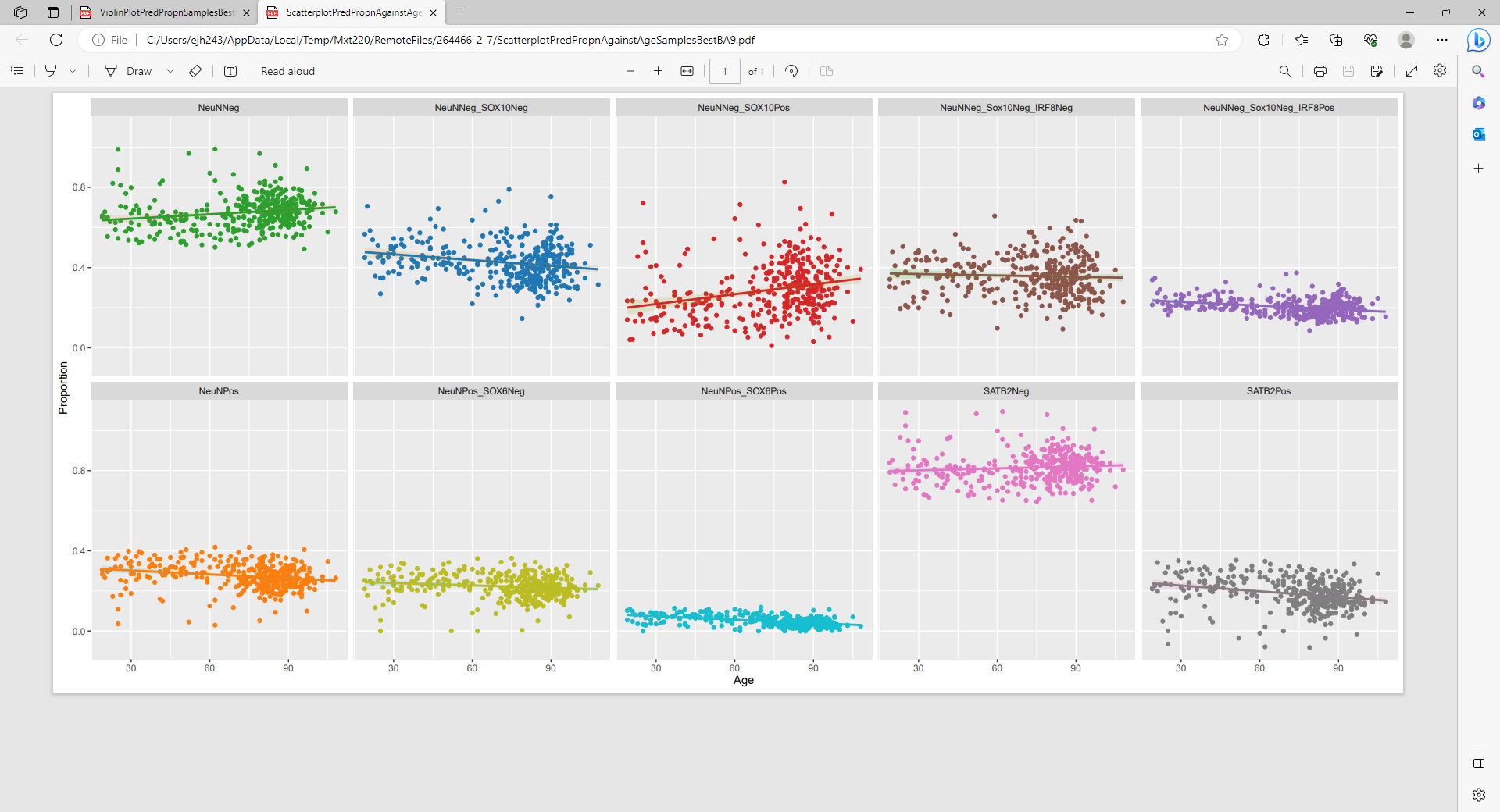

**Supplementary Figure 23.** Scatterplots plots of the of proportion of brain cell types estimated from DNA methylation data generated from the prefrontal cortex (y-axis) against age (years; x-axis) across 377 adult prefrontal cortex samples from the Exeter dataset. Each panel represents a different cell type, estimated using the optimal reference panel for that cell type (**Supplementary Table 8**).

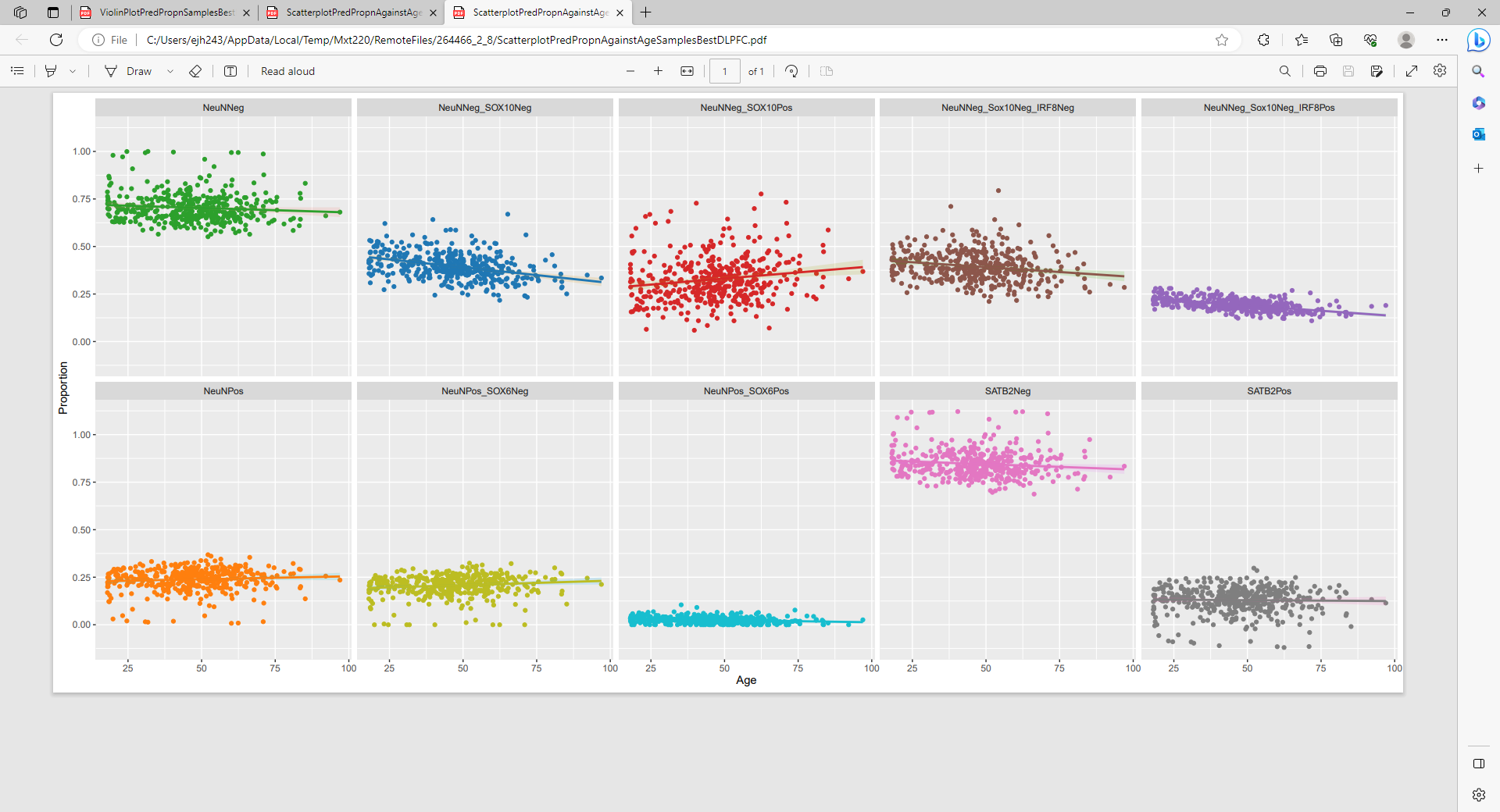
 **Supplementary Figure 24.** Scatterplots plots of the of proportion of brain cell types estimated from DNA methylation data generated from the prefrontal cortex (y-axis) against age (years; x-axis) across 415 adult prefrontal cortex samples from the Jaffe dataset. Each panel represents a different cell type, estimated using the optimal reference panel for that cell type (**Supplementary Table 8**).
